# Identifying maximally informative signal-aware representations of single-cell data using the Information Bottleneck

**DOI:** 10.1101/2024.05.22.595292

**Authors:** Serafima Dubnov, Zoe Piran, Amit Alper, Adi Yefroimsky, Hermona Soreq, Mor Nitzan

**Affiliations:** The Edmond & Lily Safra Center for Brain Sciences, The Hebrew University of Jerusalem, Jerusalem, Israel; The Alexander Silberman Institute of Life Sciences, The Hebrew University of Jerusalem, Jerusalem, Israel; School of Computer Science and Engineering, The Hebrew University, Jerusalem, Israel; Racah Institute of Physics, The Hebrew University, Jerusalem, Israel; Faculty of Medicine, The Hebrew University, Jerusalem, Israel

**Keywords:** Single-cell RNA-seq, Information Bottleneck, gene clustering

## Abstract

Rapid advancements in single-cell RNA-sequencing (scRNA-seq) technologies revealed the richness of myriad attributes encompassing cell identity. However, the complexity of the data hinders tasks focusing on a specific biological signal. To address this challenge, we introduce bioIB, a framework based on the Information Bottleneck method, designed to extract an interpretable compressed representation of scRNA-seq data, optimally-informative with respect to a desired biological signal, such as developmental stage or disease state. Provided with cellular labels representing the signal of interest, bioIB generates weighted gene clusters, termed metagenes, that compress the data, while maximizing signal-specific information. Following the Information Bottleneck principle, bioIB identifies an optimal trade-off between data compression and retaining target information. Further, bioIB provides the hierarchical structure of the metagenes, revealing the interconnections between the corresponding biological processes and cellular populations, such as the developmental hierarchy of hematopoietic cell types. We showcase bioIB’s applicability to diverse biological contexts, including Alzheimer’s Disease, epithelial-to-mesenchymal transition, immune development and hematopoiesis, demonstrating that the compressed representations capture signal-associated molecular pathways and expose cellular subpopulations with prominent phenotypes such as transition states and disease association.

## Introduction

Cellular gene expression profiles encapsulate a wealth of information regarding a cell’s identity, defined by a variety of biological factors, such as cell type, disease state, and developmental stage. Single-cell RNA-sequencing (scRNA-seq) technologies, quantifying gene expression levels at single-cell resolution, are invaluable for revealing these facets, allowing to study the different factors encompassing a cell’s identity^1^. However, exposing such factors poses a computational challenge due to the complexity and high dimensionality of scRNA-seq. While datasets typically comprise thousands of gene profiles across thousands to hundreds of thousands of cells, any reduction in dimensionality will in general result in loss of information^2^. Specifically, when aiming to uncover factors associated with a specific biological signal (e.g. gene programs associated with disease progression), the challenge can be framed as a trade-off between reducing the complexity of the data (compression) while retaining as much relevant information as possible regarding the signal of interest. The Information bottleneck (IB) theory^3^ allows us to reason mathematically about this trade-off. Given a dataset (e.g. scRNA-seq measurements) and a variable of interest encoded in the data (e.g. healthy vs. disease samples), IB provides a reduced data representation which is maximally informative about the variable of interest^3,4^. Since it was first introduced, IB has been successfully applied in diverse fields, such as text clustering^5^, image analysis^6,7^, language processing^8^, neuroscience^9^ and computational biology^10–12^.

Here, we present bioIB, a single-cell tailored method based on the IB algorithm, providing a compressed, signal-specific representation of single-cell data. The compressed representation is given by metagenes, which are probabilistic clusters of genes. The probabilistic construction preserves gene-level interpretability, allowing biological characterization of each metagene.

Previous approaches for extracting gene signatures from single-cell data include unsupervised dimensionality reduction methods, such as NMF^13^and LDVAE^14^, tools supervised by prior knowledge of signal-specific molecular pathways, marker genes and gene interactions, such as f-scLVM^15^, net-NMFsc^16^, Spectra^17^, and label-aware techniques for group-specific signature detection, such as scGeneFit^18^ and scANVI^19^. By considering the trade-off between compression and relevant information, bioIB differs from the above methods in several aspects (Table 1). Key unique aspects of bioIB include its simultaneous ability to extract gene signatures specific to a signal of interest, its independence from prior biological knowledge, and its flexibility in the number of extracted signatures or metagenes. In addition to achieving optimal signal-aware clustering of genes via metagenes, bioIB stands out from other gene program discovery tools by providing a hierarchy of metagenes, reflecting the inherent data structure relative to the signal of interest. The bioIB hierarchy facilitates the interpretation of metagenes, elucidating their significance in distinguishing between biological labels and revealing their interrelations with one another and the underlying cellular populations.

**Table 1.** Qualitative comparison of bioIB with alternative methods for the generation of gene signatures from single-cell data.

| Method | Method output | Signal-specific | Control over the number of gene signatures | Option of hierarchical representation of gene factors |
| --- | --- | --- | --- | --- |
| bioIB | Probabilistic gene signatures | ✓ Yes<br>• Generating signal- or condition- associated gene signatures (Figures 2 – 6) | ✓ Yes<br>• Elucidating intermediate and transition signatures (Figures 2,3)<br>• Separating condition-specific cellular subpopulations (Figures 4,5) | ✓ Yes<br>• Revealing the biological interconnections between the gene factors, as well as interconnections between the underlying cellular subpopulations (Figures 4-6) |
| scGeneFit | Gene markers | ✓ Yes | ✗ No | ✗ No |
| scANVI | Condition-specific gene rankings | ✓ Yes | ✗ No | ✗ No |
| NMF | Weighted gene signatures | ✗ No | ✓ Yes | ✗ No |
| LDVAE | Weighted gene signatures | ✗ No | ✓ Yes | ✗ No |

We demonstrate that metagenes generated by bioIB are biologically meaningful, capturing molecular pathways differentially activated between signal-specific cell groups. First, using a scRNA-seq dataset of neurons with and without Alzheimer’s Disease (AD) associated neurofibrillary tangles^20^, we show that bioIB metagenes capture relevant molecular pathways enriched in each group and in the intermediate transcriptomic state, elucidating more signal-related genes compared to competing methods and clustering them in agreement with known biological pathways. We further demonstrate that bioIB metagenes capture cells in the transition state in the context of epithelial-to-mesenchymal transition (EMT) scRNA-seq dataset^21^. Next, applying bioIB to an atlas of differentiating macrophages^22^, and using either organ-of-origin or developmental stage as signals of interest, we show that bioIB extracts distinct, signal-specific metagene hierarchies and associated biological processes. We also demonstrate how bioIB can be used to identify a cellular subpopulation of disease-associated astrocytes in a single nucleus RNA-seq (snRNA-seq) dataset^23^ from murine Alzheimer’s Disease models. Finally, we showcase that bioIB metagene hierarchy for a dataset of differentiating hematopoietic cell types^24^ reflects the developmental hierarchy of the corresponding cellular populations. bioIB is available as an open-source software package, along with documentation and tutorials (https://github.com/nitzanlab/bioIB).

## Results

### bioIB elucidates signal-specific metagenes and their structure

The bioIB representation is computed for a given dataset and signal of interest, provided as cell labels. The representation is composed of metagenes which are probabilistic aggregation of the genes into clusters, representing the major patterns of gene expression variation underlying the labeled signal.

The input to bioIB includes a count matrix *D* ∈ *R*^*N*×*G*^ of N cells by G genes, and a vector of cell labels related to the signal of interest, *S* ∈ *R*^*N*×1^, where for example, each cell is labeled as sampled from either a healthy or diseased population (Methods; Figure 1A). This input is used to estimate the distributions required for the IB algorithm. We thus define three categorical random variables, *C* ~*Cat*({*c*_1_, …, *c*_*N*_}), *X* ~*Cat*({*x*_1_, …, *x*_*G*_}), *Y* ~*Cat*({*y*_1_, …, *y*_*K*_}), respectively representing the *N* cells, *G* genes and *K* cell states of interest. Normalizing the input matrix *D* by the total number of counts, we obtain a joint probability distribution *p*(*c, x*). Next, summing *p*(*c, x*) across the cells, we obtain *p*(*x*), such that an entry [*p*(*x*)]_*i*_ represents the marginal probability of sampling the transcript of gene *x*_*i*_.

**Figure 1.**
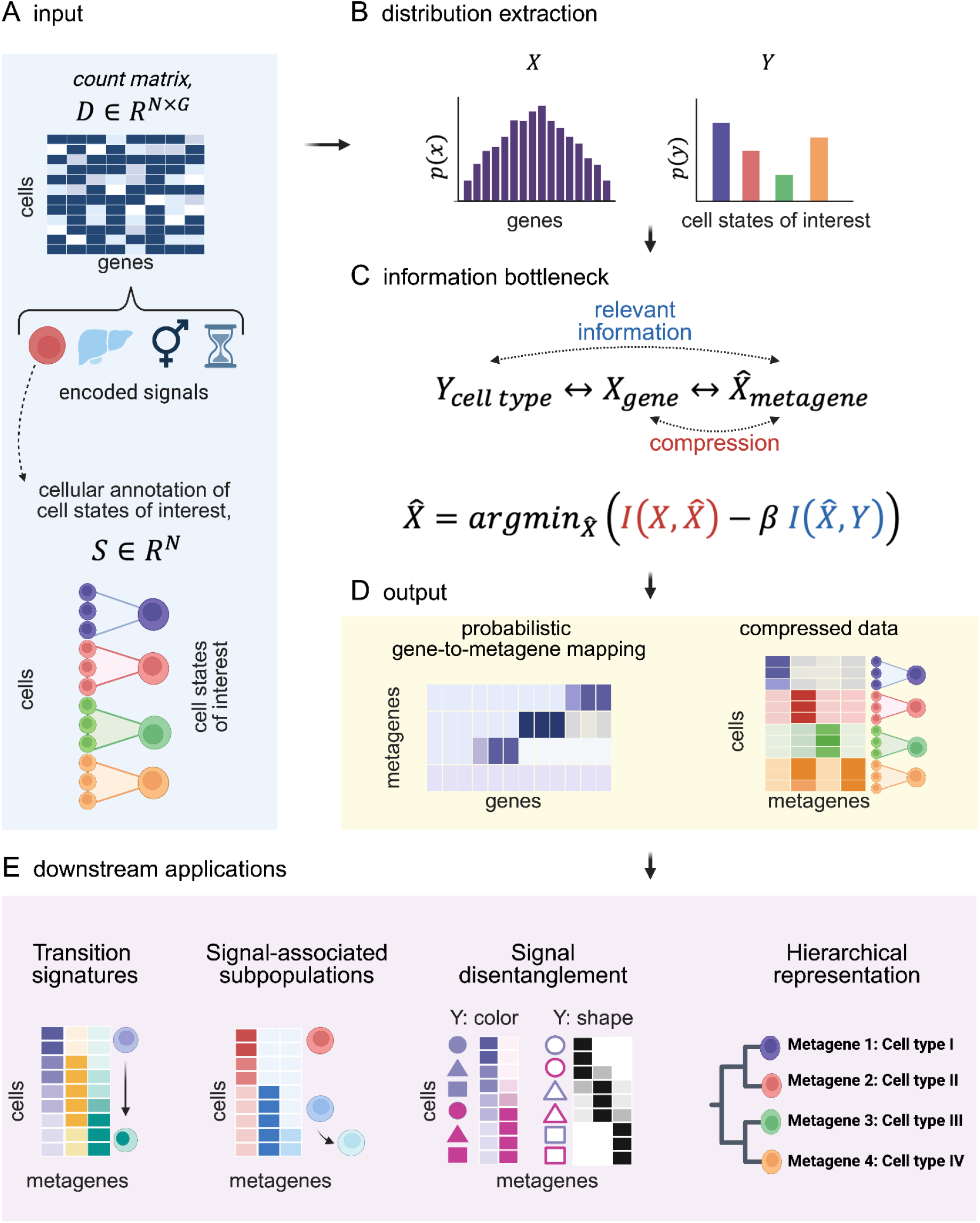
Elucidating meaningful, signal-specific metagenes using bioIB. A-D) The bioIB pipeline. A) Input; bioIB takes as input a gene count matrix and a cellular annotation vector, labeling every cell with a state, representing the signal of interest. For example, if the signal of interest is cell type, these labels annotate every cell with the corresponding cell type. B) Distributions extraction; The provided count matrix and the cellular annotation vector are used to estimate the distributions of the random variables representing the genes (*X*) and the cell states of interest (*Y*). C) Information Bottleneck (IB); The probabilities obtained in (B) are used as input for the IB algorithm, which yields the optimal mapping of genes to metagenes, by optimizing the trade-off between compression, linking genes (*X*) and metagenes 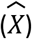, and relevant information, linking metagenes 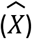 and the cell states of interest (*Y*). This is achieved by optimizing for 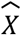 that minimizes the mutual information with the input genes 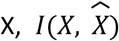, while maximizing the mutual information with 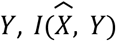. D) Output; The output of bioIB is a probabilistic mapping between genes and metagenes, scoring all the genes measured in the input matrix by their contribution to each metagene. bioIB also provides a cell-to-metagene compressed representation of the input matrix, summarizing the expression of metagenes in single cells. E) Possible downstream applications of the compressed data achieved by bioIB: elucidating transition signatures, identifying signal-associated cellular subpopulations with distinct transcriptional profiles, disentangling distinct label-specific representations, and characterizing the hierarchical interconnections between metagenes and the corresponding cell types. Figure was created with BioRender.com.

Using Bayes theorem, we obtain the conditional probability:

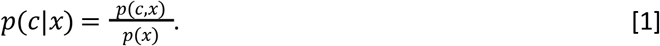

Here, an entry [*p*(*c*|*x*)]_*ij*_ represents the probability that a randomly sampled cell from *D* is the cell *c*_*i*_, given that it expresses gene *x*_*j*_. The provided cellular annotation vector *S* ∈ *R*^*N*×1^ allows us to define the conditional distribution of *Y* (representing the *K* cell states of interest) given that we observed a cell in *D*. By definition *p*(*y*|*c*) is an indicator function, defined by *S*, namely, for a cell *c*_*i*_, *p*(*y*|*c* _*i*_) = 1 if *S*_*i*_ = *y* and zero otherwise:

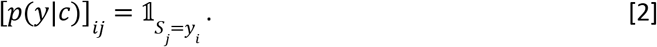

At last, we can obtain the conditional distribution of cell states of interest given that we observed a certain gene in *D*:

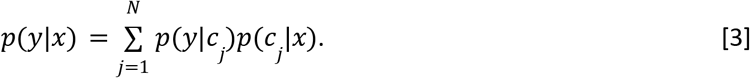

The conditional probability matrix of cell states given the genes 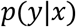 and the gene probability vector *p*(*x*) are used as input to the core of the bioIB method, the IB algorithm.

The IB yields the optimal probabilistic mapping, 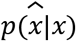 from the genes’ random variable, *X*, to the categorical random variable representing the metagenes 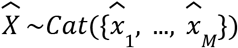, (for |*M*| <= |*G*|). The mapping is optimal with respect to the tradeoff between compression and information about the signal of interest *Y* according to a given threshold parameter β (Figure 1C). This is achieved by optimizing for 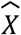 that minimizes the mutual information with the input genes 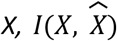, while maximizing the mutual information with 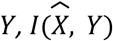 (Methods; Figure 1C):

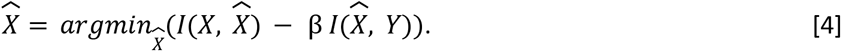

The resulting metagenes are probabilistic clusters of genes capturing the shared expression patterns amongst cell states relative to *Y* (Figure 1D). The metagenes are defined by two probabilistic matrices, one linking metagenes to genes 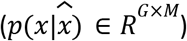 and another - linking metagenes to cell states of interest 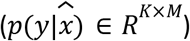. In the flat clustering mode, bioIB generates *M* metagenes, where *M* is defined by the user (Methods). Additionally, bioIB can obtain a hierarchy of metagenes by gradually decreasing β through a reverse-annealing process^4^ (Methods). In the hierarchical mode, the number of metagenes *M* is roughly determined by the threshold parameter β, ranging from the original representation (no compression,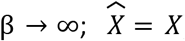) to full compression to a single cluster (β = 0). The probabilistic output mapping, 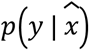, reflects the amount of information each metagene holds regarding the different labels, whereas the hierarchical structure reveals the interdependence between the metagenes, and the underlying cellular populations they correspond to (Figure 1E). As an illustrative example, we construct a toy dataset composed of cells belonging to one of two cell types, which act as the signal of interest Y (Supplementary Figure 1A-D). The bioIB hierarchy is revealed by plotting the conditional probabilities 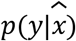 of a particular label given every metagene, across β values that define the compression level (Supplementary Figure 1C-D). The hierarchical structure reflects the interconnections among the metagenes and the specified cell types of interest (*Y*), while the bifurcation order is dictated by the informativity of the generated metagenes relative to *Y*. bioIB can also capture the relationships between related cell types, defined as distinct labels of interest (*Y)*. Given a toy model with four related cell types, bioIB hierarchy reflects the two distinct pairs of linked cell types by two branches. Further splits correspond to higher-resolution separation to different cell types, eventually resulting in cell type-specific metagenes (Supplementary Figure 1E-G). Progressing to simulated data that more realistically reflects the characteristics of scRNA-seq^25^, we show that bioIB outperforms competing methods (including scGeneFit^18^, scANVI^19^, NMF^13^ and LDVAE^14^) in identifying underlying signal-specific genes (Supplementary Figure 2). Furthermore, bioIB is robust to batch effects, class imbalance, erroneous cellular annotations, and cell subsampling (Supplementary Figures 3-6).

### bioIB elucidates a spectrum of gene programs underlying the gradual development of the pathological phenotype in Alzheimer’s Disease neurons

Both clinical^26^ and pathological^27,28^ manifestations of Alzheimer’s Disease (AD) suggest that it is a continuum with a gradual development of the pathological phenotype. Here we show that by tuning the number of metagenes, bioIB reveals a spectrum of gene signatures underlying the gradual transformation associated with Alzheimer’s Disease (AD). We applied bioIB to a scRNA-seq expression profiles of excitatory neurons with and without the neurofibrillary tangles (NFT)^20^ to elucidate molecular pathways underlying neuronal vulnerability in AD. Given the cellular labels indicating the presence or absence of tau pathology (NFT-bearing vs. NFT-free, respectively; Figure 2A), the three metagenes generated by bioIB (Methods) revealed a gradual shift in expression levels with metagenes 0 and 2 overrepresented in NFT-free and NFT-bearing neurons, respectively, and metagene 1 representing an intermediate state signature between the two populations (Figure 2B, Supplementary Tables 1,2). By dividing cells to metagene-associated clusters based on their relative metagene expression (Supplementary Figure 7A; Methods), and assessing the phenotype progression stage using the NFT-associated gene markers^20^ (Figure 2C, Supplementary Figure 7B), we found that indeed, the NFT-linked marker genes gradually increased in expression from metagene 0, associated with NFT-free neurons, through the ‘intermediate state’ metagene 1, to metagene 2, associated with NFT-bearing neurons (Figure 2C, Supplementary Figure 7B). Next, the direct link between bioIB’s metagenes to genes (Figure 2D) allowed us to interpret the biological identity of each metagene (Figure 2E; Methods, Supplementary Table 3). Metagene 0, associated with the NFT-free cells, was enriched for axon guidance, an essential pathway of neuronal homeostasis^29^. Metagene 2, associated with NFT-bearing cells, was represented by multiple genes linked to Alzheimer’s Disease progression^30–33^ enriched in synaptic plasticity and neurotransmitter secretion, in agreement with previous findings^20^ (Figure 2E). Finally, metagene 1, representing the intermediate state, is enriched for oxidative phosphorylation, reported to be damaged at the early stages of the disease^34^ (Figure 2E).

**Figure 2.**
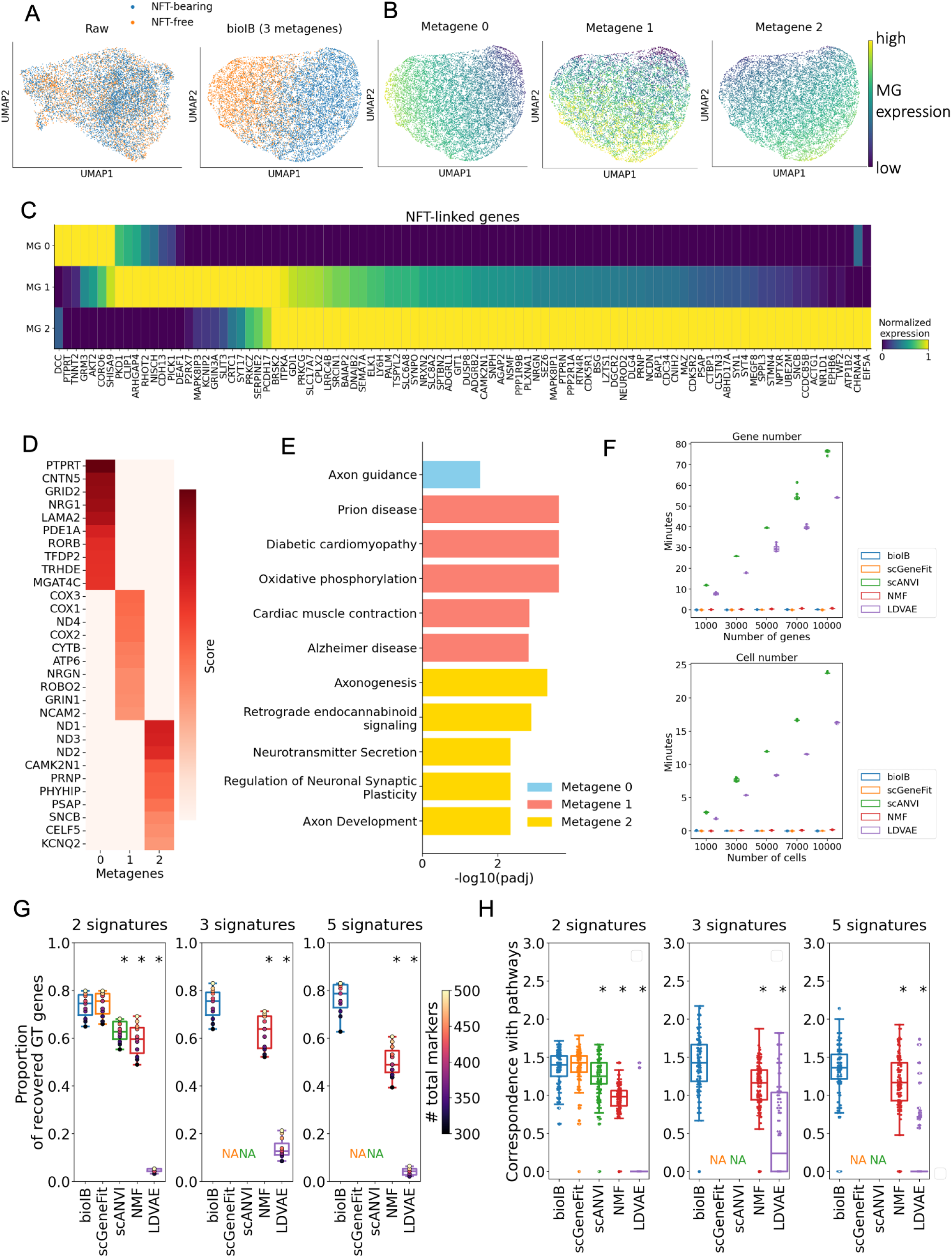
bioIB elucidates a spectrum of gene programs underlying the development of the pathological cellular phenotype in Alzheimer’s Disease neurons. A) UMAP representation of the original data^20^ (left) and of the bioIB compressed data (right), colored by the input labels indicating the presence of NFT pathology. B) UMAP representation of the bioIB compressed data, colored by the expression levels of the resulting metagenes (Methods). C) Heatmap featuring normalized expression of NFT-associated gene markers (as previously defined^20^) in cells clustered by relatively maximized metagene expression (Methods). D) Heatmap featuring the top 10 genes representing each of the metagenes and their corresponding probabilistic metagene-to-gene mapping. E) Barplot showing the top significant pathways (GO Biological Process, KEGG pathways) enriched within the top 100 markers of each metagene. F) CPU Runtime as a function of the number of genes (given 10,955 cells; top) and number of cells (given 3,000 highly variable genes; bottom) for bioIB, scGeneFit, scANVI, NMF and LDVAE. The experiment was repeated *n* = 10 times. G) Fraction of ground-truth informative genes, shown in (C) (Methods) recovered within top 300-500 gene markers of two (left), three (middle) and five (right) gene signatures generated by bioIB, scGeneFit, scANVI, NMF and LDVAE. Statistical significance was assessed using the WIlcoxon signed rank test (non-parametric), with * indicating p < 0.01, in comparison to the bioIB scores. H) Correspondence between the division of genes to signatures and ground-truth pathways for two (left), three (middle) and five (right) gene signatures generated by bioIB, scGeneFit, scANVI, NMF and LDVAE. For each method, the correspondence scores were normalized to the scores of shuffled signatures, per pathway (Methods). Statistical significance was assessed using the Mann–Whitney U-test (non-parametric), with * indicating p < 0.01, in comparison to the bioIB correspondence scores. In box plots middle line, median; box boundary, interquartile range (IQR); whiskers, 1.5*IQR; gray dots, points beyond the minimum or maximum whisker. \**MG – metagene*.

We defined a set of benchmark tasks aimed to assess the biological interpretability of outputs of different methods by quantifying their similarity to the molecular signatures of neuronal vulnerability^20^ (Supplementary Table 4, Methods). First, we compared the fraction of recovered informative genes (characterized as part of the neuronal vulnerability signatures) captured within the top (300/500) markers of the produced metagenes, or gene factors. bioIB outperforms competing methods in recovering informative genes given three and five gene signatures that expose the intermediate condition, and performs similarly to scGeneFit while outperforming other baselines given two gene signatures (one signature per condition, Figure 2G). We additionally evaluated the correspondence between produced metagenes or factors and previous division of genes to biological pathways^20^ (Supplementary Table 4). bioIB outperforms competing methods in informative pathway recovery given three and five gene signatures, and performs similarly to scGeneFit while outperforming other baselines given two gene signatures (Figure 2H, Supplementary Figure 8, Methods).

bioIB’s low runtime and moderate memory usage make it suitable for running on CPUs, even with large datasets, especially when restricted to highly variable genes, as recommended (Figure 2F, Supplementary Figure 9, Supplementary Table 5). At last, bioIB is robust to initialization parameters and noisy data (Supplementary Figures 10, 11).

### bioIB identifies cells at the transition state between epithelial and mesenchymal phenotypes

Biological signals often represent gradual transition processes, with the cellular labels signifying their correspondence to the end-point phenotypes. In this scenario, apart from the state-specific binary markers, the transition genes expressed along the trajectory are of particular interest. We studied this setting in the context of the epithelial-to-mesenchymal transition (EMT) by applying bioIB to the analysis of a scRNA-seq data from primary human mammary epithelial cells^21^. Given the cellular annotation (epithelial or mesenchymal), we used bioIB to generate three metagenes, with two metagenes enriched in either mesenchymal or epithelial states (metagenes 0 and 1, respectively), and one metagene enriched in a transition stage (metagene 2; Figure 3A; Supplementary Tables 6,7). Notably, the state-specific metagenes exhibited a gradual expression change, correlated with the EMT transition. In particular, the expression of metagene 0 monotonically decreases (increases) with mean marker expression of epithelial (mesenchymal) marker genes (Spearman correlation coefficients −0.43 and 0.7, respectively) (Figure 3B). On the contrary, the expression of metagene 1 monotonically increases (decreases) with the epithelial (mesenchymal) marker expression (Spearman correlation coefficients 0.54 and −0.58, respectively) (Figure 3B). Metagene 2 exhibited weaker correlation with markers of both phenotypes (Spearman correlation coefficients 0.22 and −0.13 for epithelial and mesenchymal markers, respectively). Consequently, cells maximizing metagene 0 (metagene 1) feature a differentiated mesenchymal (epithelial) phenotype, whereas cells maximizing metagene 2 represent a transition state between the two phenotypes (Figure 3C), and express intermediate levels of epithelial and mesenchymal marker genes (Figure 3D). Furthermore, the transition (metagene 2) signature is enriched for categories related to p53 pathway, Myc signaling and translation (Figure 3E, Methods), in agreement with previous findings^21^.

**Figure 3.**
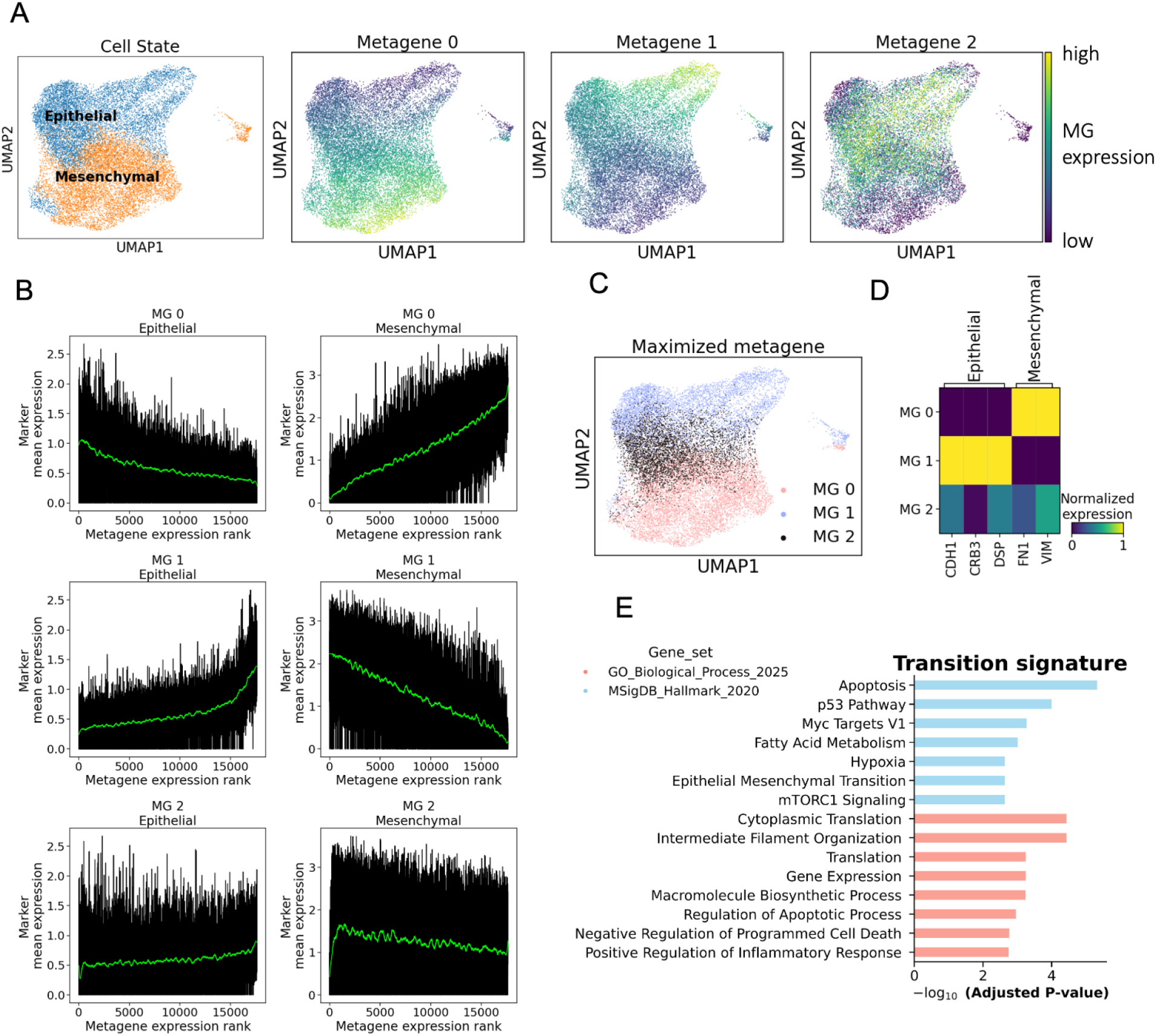
bioIB identifies cell states along the epithelial to mesenchymal transition. A) UMAP representation of the original data, colored by the input labels of epithelial and mesenchymal phenotypes (left), and by the relative expression of bioIB metagenes (right). B) The mean expression level of epithelial (left column) and mesenchymal (right column) marker genes as a function of the expression level ranks of metagene 0 (top row), metagene 1 (middle row) and metagene 2 (bottom row), per cell. C) UMAP representation of the original data colored by the relatively maximized metagene. D) Heatmap showing the relative expression levels of epithelial and mesenchymal markers in three metagene-associated cellular populations shown in (C). E) Barplot with the enriched GO Biological Processes and MSigDB Hallmark pathways within the top 100 markers of metagene 2, representing the transition signature. \**MG – metagene*.

### bioIB extracts distinct molecular signatures in macrophages for developmental stage and organ residence across development

Gene expression data in scRNA-seq experiments contain signatures associated with multiple overlapping biological signals or conditions. How can we identify gene signatures associated with a specific source of heterogeneity in the data? We demonstrate bioIB’s approach to this challenge in the context of a scRNA-seq atlas of the developing immune system, which contains cells from multiple organs spanning weeks 4 to 17 after conception^22^ (Figure 4A). We focused on the macrophages population, due to the variability of their gene expression across organs and throughout the gestation stages, with specific subpopulations, differentially abundant both between different organs and across development^22^ (Supplementary Figure 12). Here we demonstrate that bioIB metagenes are associated with specific macrophage subpopulations, and that the bioIB hierarchy reveals their interconnections with respect to the signal of interest.

**Figure 4.**
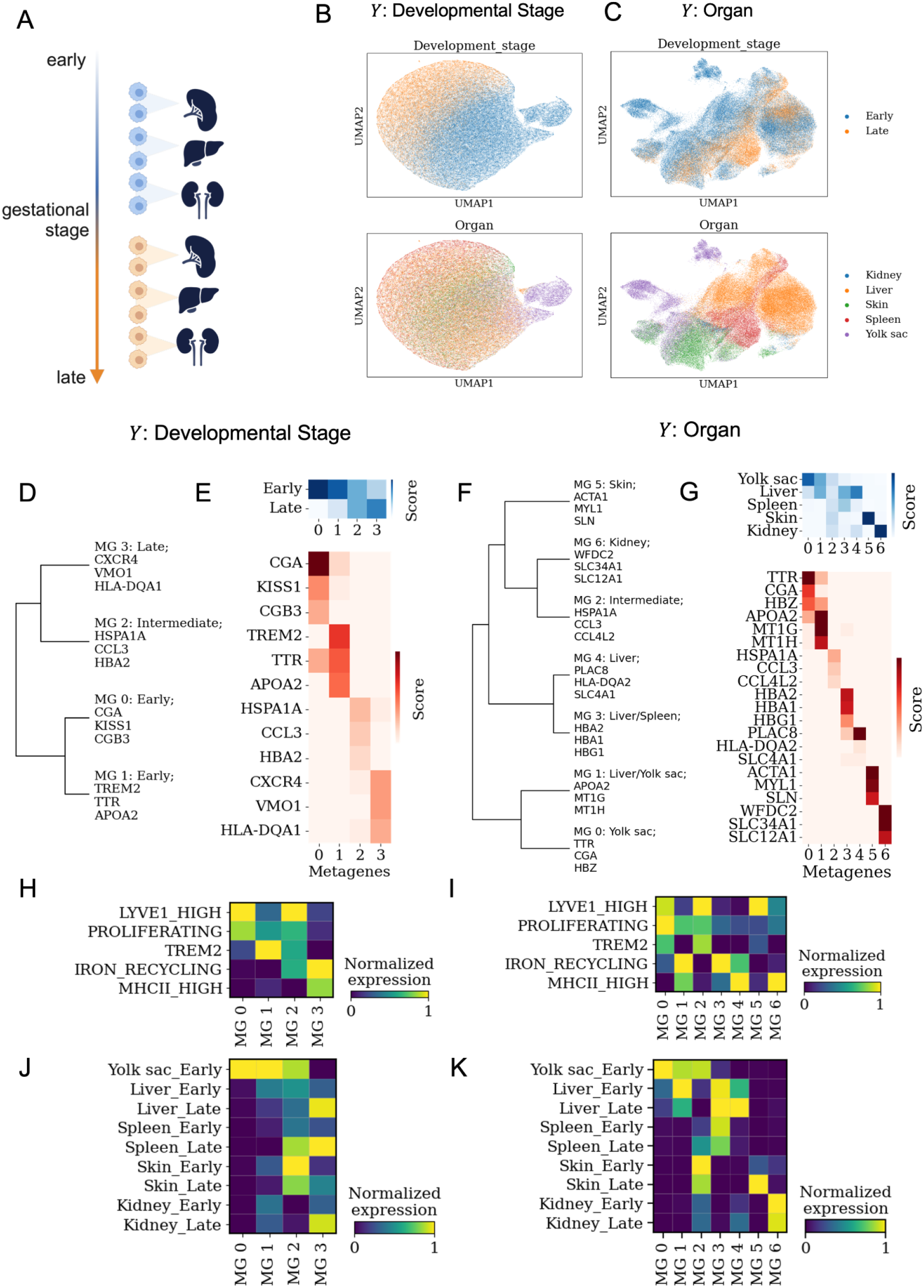
bioIB extracts distinct molecular signatures underlying the signals related to developmental stage and organ-of-origin in developing macrophages. A) Schematic representation of the analyzed scRNA-seq data^22^ of macrophages from 5 distinct organs (kidney, liver, skin, spleen, yolk sac) and 11 gestational weeks (4, 7-12, 14-17). Figure created with Biorender.com. B,C) UMAP representation of the data compressed by bioIB with Y set either to (B) developmental stage (Early: < 14 weeks; Late: >= 14 weeks) or (C) organ-of-origin, colored by the developmental stage (top) or by organ-of-origin (bottom). D,F) Metagene hierarchy inferred from bioIB with Y set either to (D) developmental stage or (F) organ-of-origin. Each metagene is labeled with the associated cell group(s) of interest and three top representative genes (Methods). E,G) Heatmaps showing the probabilistic mappings between bioIB metagenes and cell groups of interest (top) and genes (bottom) with Y set either to (E) developmental stage or (G) organ-of-origin. H,I) Heatmaps representing the relative expression of bioIB metagenes generated with *Y* set to (H) developmental stage or (I) organ-of-origin, in macrophage subpopulations defined by the original analysis^22^. J,K) Heatmaps representing the relative expression of bioIB metagenes generated with *Y* set to (J) developmental stage or (K) organ-of-origin, in cellular clusters divided by organ-of-origin and developmental stage. \**MG – metagene*.

Using the hierarchical mode of bioIB via a reverse-annealing process (Methods), we gradually compress the data, subsequently merging the metagenes carrying similar biological information about the selected signal of interest and thus exposing a signal-specific hierarchy of gene programs. The bioIB hierarchy is based on the probabilistic mapping between cellular labels and metagenes across a range of β values, representing the clustering resolution, or the number of metagenes (Methods).

We first applied bioIB with *Y*, the signal of interest, set to be the developmental stage, after aggregating cells, each assigned either ‘Early’ (8-12 gestational weeks) or `Late’ (>14 gestational weeks) label. The resulting bioIB representation comprised four macrophage-specific metagenes (Methods, Supplementary Figure 13), enhancing the target signal of the developmental stage (Figure 4B; Supplementary Tables 8,9). These four metagenes were organized into two branches: two metagenes (0, 1) associated with the early stage, and two metagenes (2, 3) associated with the intermediate stage (2) and the late stage (3) (Figure 4D,E; Supplementary Figure 13, Methods). The stage-specific metagenes (0,1,3) were upregulated in the relevant macrophage subpopulations (Figure 4H). The early stage-associated metagenes 0 and 1 are enriched in LYVE-high, proliferating macrophages and TREM2-positive macrophages, respectively, while the late stage-associated metagene 3 is enriched in iron-recycling and MHCII-high macrophages, in agreement with previous findings^22^. The intermediate metagene 2 was found to be enriched both in all early stage-associated subpopulations (LYVE-high, proliferating and TREM2-positive macrophages), as well as in the late stage-associated iron recycling macrophages. Finally, comparing the metagene expression between cellular groups divided both by the developmental stage and the organ-of-origin supported the specificity of bioIB metagenes to the selected signal of interest (Figure 4J). Indeed, metagenes 1 and 3 are respectively overrepresented in early and late cells of all organs, revealing stage-specific gene programs, common to multiple organs. Furthermore, metagene 0 represents a distinct signature of early stage-associated genes, specifically increased in yolk sac. Thus, bioIB can both capture the dominant signal-associated transcriptional patterns shared across cells and identify subpopulations that deviate from these common patterns.

When *Y* is set to be the organ-of-origin (Figure 4C), the bioIB hierarchy exposes both the organ-specific and the shared transcriptional programs, revealing the macrophage subpopulations with similar phenotypes across different organs (Figure 4F,G; Supplementary Tables 10, 11). The yolk sac branch (metagenes 0, 1) differentiates between a yolk sac-specific signature enriched in macrophages from LIVE-high, proliferating and TREM2-positive populations (metagene 0), and an additional gene program shared between the yolk sac and the liver macrophages, enriched in proliferating and iron-recycling populations, reflecting the shared hematopoietic properties of the yolk sac and the liver^35^ (metagene 1) (Figure 4F,G,I,K). In parallel, while metagene 4 represents a liver-specific signature, metagene 3 elucidates a transcriptional signature shared between the liver and the spleen, also enriched in iron-recycling macrophages, in agreement with previous findings^22^ (Figure 4F,G,I,K). Finally, the organ-specific metagenes elucidate the genes associated with particular organs or organ groups, and appear to be generally common across developmental stages (Figure 4K).

### bioIB metagenes identify Alzheimer’s Disease associated astrocytes

A key challenge in scRNA-seq analysis is to identify specific cellular subpopulations affected by a certain condition, such as disease. The standard pipeline, commonly implemented for this task, involves unsupervised clustering of cells, which exposes the downstream analysis to clustering-related bias^36^. BioIB can overcome such limitations and detect disease-associated cells within a heterogeneous cellular population, which we demonstrate in the context of Alzheimer’s disease (AD) - associated astrocytes. To do so, we re-analyzed single-nucleus RNA-seq measurements of astrocytes from an AD mouse model and wild-type (WT) mice^23^ (Figure 5A).

**Figure 5.**
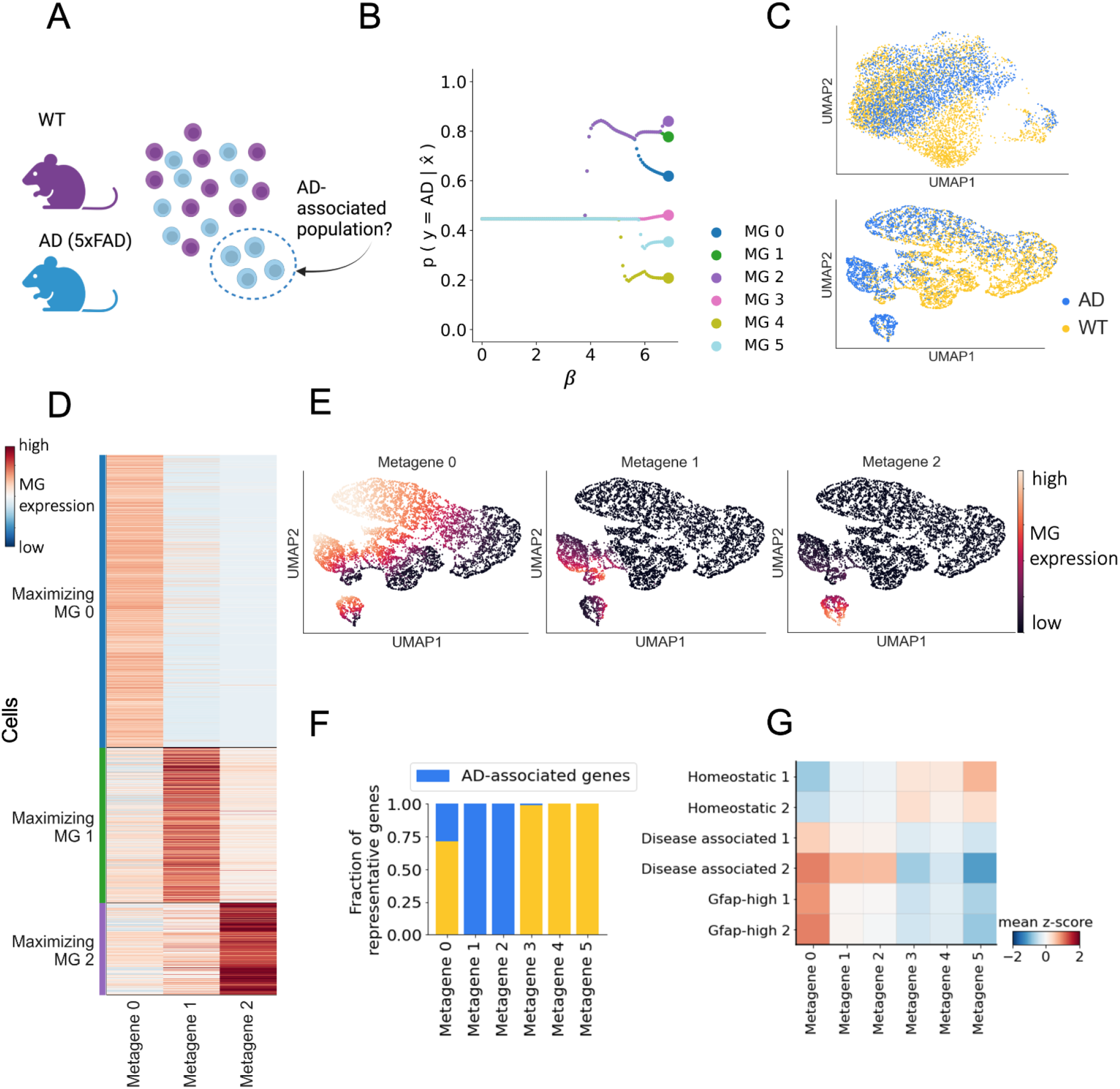
bioIB metagenes reveal AD-associated astrocytes. A) Schematic representation of the snRNA-seq dataset of astrocytes, derived from a murine model of AD ^23^. The data was analyzed using bioIB with *Y* set to genotype (WT/AD), which resulted in identification of a specific subpopulation of disease-associated astrocytes. Figure created with Biorender.com. B) BioIB metagene hierarchy produced given the preprocessed snRNA-seq data, relating to the AD group. The defined metagenes exhibit differential expression patterns between AD and WT, with metagenes 0, 1 and 2 overexpressed in AD cells (Fold change increase in metagenes 0, 1, 2: 1.9, 3.9, 6, respectively), a neutral metagene 3 (Fold change increase in metagene 3 = 0.91), and metagenes 4 and 5 overexpressed in WT cells (Fold change increase in metagenes 4, 5: 0.3, 0.66, respectively; Supplementary Table 15). C) UMAP representation of the original data (left) and of the bioIB compressed data (right). D) Heatmap showing scaled expression of metagenes 0,1,2 in individual cells of AD genotype, sorted by maximal normalized metagene expression. E) UMAPs of the bioIB-compressed data, colored by the expression of AD-associated metagenes 0,1,2. F) Fractions of representative genes of metagenes 0-5 that were found to be differentially expressed in at least 7 studies in the meta-analysis of the AD-associated transcriptome^48^ (Methods). G) Heatmap of scaled expression values of six bioIB metagenes in six transcriptional clusters of astrocytes, defined in ref^23^. \**MG – metagene*.

BioIB analysis with the signal of interest set as the genotype (AD/WT) resulted in a hierarchy of six metagenes (Supplementary Tables 13,14) capturing informative transcriptomic signatures differentiating between AD and WT cells (Figure 5B,C, Supplementary Figure 16A). Furthermore, bioIB metagenes captured a higher-resolution structure within the data; the main branch of metagenes associated with AD genotype is composed of metagenes 0,1,2, each associated in turn with a distinct subpopulation of AD astrocytes (Figure 5D,E). To interpret their biological identities, we extracted a set of representative genes for each metagene (Methods). Metagene 0, whose representative gene set includes genes involved in morphology regulation (*GFAP, THY1, VIM, B2M, PSEN1*), is enriched for the cellular projection development process, consistent with general astrocyte activation^37–42^ (Supplementary Figure 16B,C). Metagenes 1 and 2 represent pathways more tightly associated with the disease:, the representative gene set of metagene 1 is enriched with immune genes^43^, such as C1QA^44^ and CTSS^45^, and metagene 2 is represented by established markers of AD pathology, *TYROBP* and SERPINA3N^46,47^. Meta-analysis of the AD-associated transcriptome^48^ revealed that metagenes 1 and 2 are the only metagenes that are exclusively represented by AD-associated genes (Figure 5F; Methods). Characterization of the WT-related metagene 5 can be found in Supplementary Figure 16D.

While metagene 0 is expressed in the majority of AD astrocytes, metagenes 1 and 2 characterize distinct cellular subpopulations among the AD cells (Figure 5D,E), which we hypothesized to correspond to disease-associated astrocytic signatures. To support our interpretation, we quantified the expression of bioIB metagenes in six astrocytic clusters defined in^23^, which included two homeostatic clusters, two GFAP-high clusters of reactive astrocytes which are not specific to the disease, and two disease-associated clusters^23^. We found that while bioIB metagene 0 is highly expressed both in disease-associated clusters and in reactive GFAP-high clusters, metagenes 1 and 2 are specifically enriched in the disease-associated cluster, most abundant in AD^23^ (Figure 5G; Supplementary Figure 16E). The two WT-associated metagenes (4,5) are correspondingly enriched in the homeostatic clusters (Figure 5G). In summary, bioIB allows to directly uncover the cellular subpopulations differentially affected by the disease.

### bioIB metagene hierarchy reflects developmental connections between hematopoietic cell types

scRNA-seq datasets expose a striking diversity of cell types and states, whose interconnections carry important biological information about cell state identity. For example, the hierarchical differentiation tree of hematopoietic stem and progenitor cells (HSPCs) reveals the phenotype and function of mature hematopoietic cells^49^. BioIB metagene hierarchy can capture the developmental hierarchical structure of cell types, as we demonstrate here for scRNA-seq data of HSPCs differentiation^24^ (Figure 6A). BioIB is applied given the cell type signal over a subset of the data containing six major hematopoietic cell types - monocytes, neutrophils, mast cells, basophils, megakaryocytes and erythroid cells. This analysis produced 11 metagenes, where each of the six cell types is uniquely characterized by at least one metagene, maximizing its expression level within that particular cell type (Figure 6B, Supplementary Tables 16,17). In addition, there are metagenes representing a transcriptional program shared by several developmentally linked cell types (Figure 6B,C; Supplementary Figure 17A). For example, metagenes 0 and 2 are specifically expressed in monocytes and neutrophils, respectively, while metagene 1 is activated in both (Figure 6B,C). The bioIB metagenes are biologically informative, uniting genes and processes characteristic of the corresponding cell types (Supplementary Figure 17B,C). Hence, metagene 0, specifically representing monocytes, features monocyte marker genes such as FABP5^24^ and WFDC17^50,51^ (Figure 6C,D), and is associated with pro-inflammatory macrophage activation, characteristic of monocytes function^52^ (Figure 6E). Similarly, metagene 2, specifically characterizing neutrophils, includes markers like ITGB21^24^ CAMP, LTF, and ELANE^53^ (Figure 6C,D) and is statistically enriched for neutrophil mediated immunity and neutrophil activation (Figure 6E).

**Figure 6.**
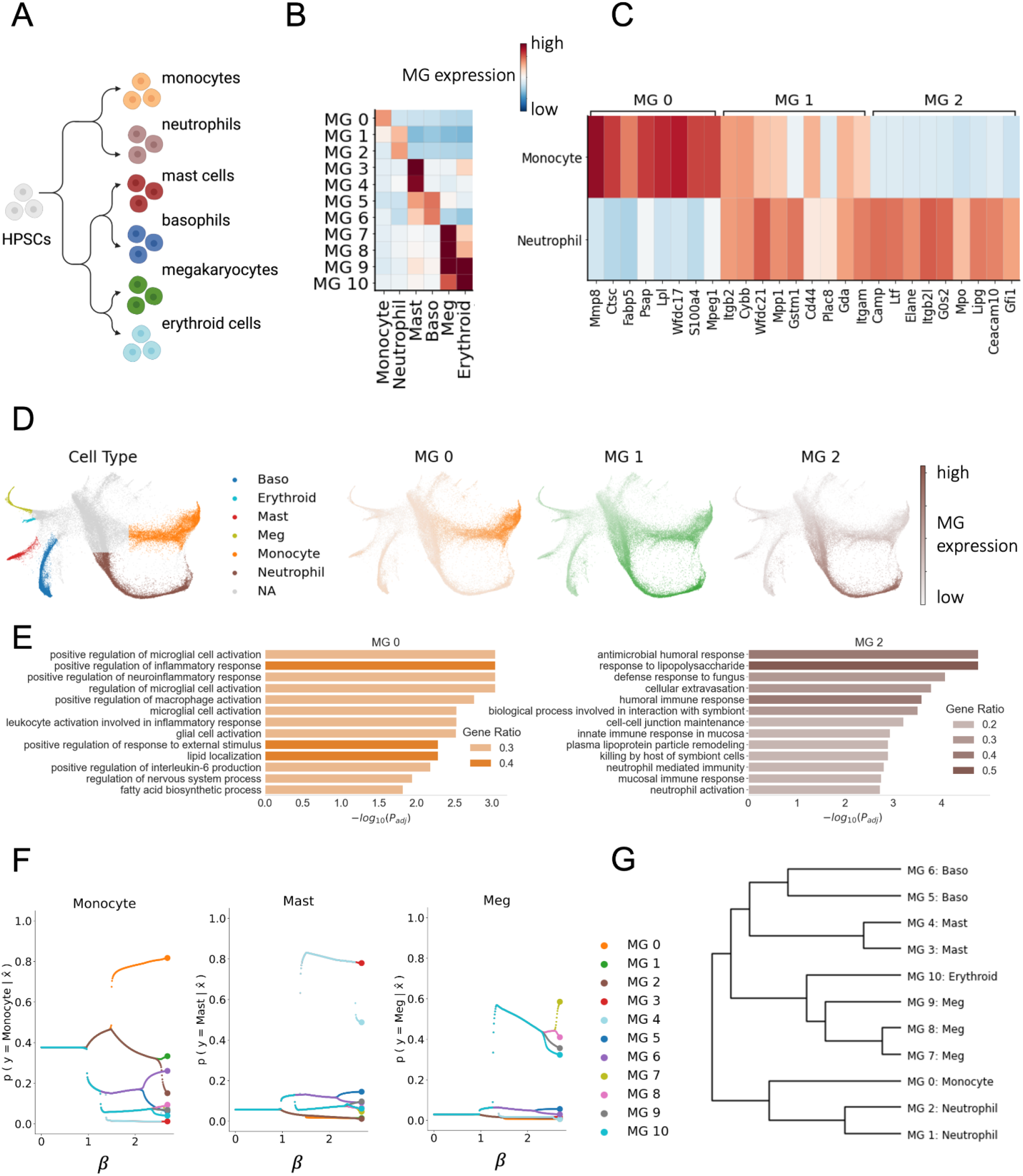
bioIB metagene hierarchy reflects the connections between the developmentally linked hematopoietic cell types. A) Schematic representation of the scRNA-seq dataset of differentiating hematopoietic cell types^24^ with their associated developmental hierarchy. Figure created with Biorender.com. B) Heatmap showing the scaled expression (z-score) of the bioIB metagenes across cell types. C) Heatmap showing scaled expression of the top representative genes of metagenes 0,1,2 across monocytes and neutrophils. Metagenes 0 and 2 are specifically expressed in monocytes and neutrophils, respectively, while metagene 1 is expressed in both. D) SPRING54 visualizations of the hematopoietic dataset (embedding as provided in^24^), colored by cell type (left panel) and by the expression of metagenes 0-2 (three panels on the right). E) Gene Ontology enrichment results showing biological process categories significantly enriched in metagene 0 (left) and 2 (right). F) Bifurcation plots of further compression of the 11 metagenes shown in (B) relative to Monocytes, Mast cells and Megakaryocytes. Metagenes characterizing developmentally linked cell types are linked in the metagene hierarchy. For example, metagene 0 representing monocytes diverges from the same branch as metagene 2, representing Neutrophils. Bifurcation plots relative to Neutrophils, Basophils and Erythroid cells are provided in Supplementary Figure 4D. G) Metagene hierarchy inferred from the bioIB reverse annealing output shown in (F) and in Supplementary Figure 4D. The cell type associated with every metagene is the one maximizing the conditional probability 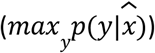 of a cell type *y* given this metagene, 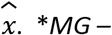 *metagene*.

The hierarchical representation of the metagenes generated by bioIB induces a hierarchy of cell types that reflects the developmental links between them (Figure 6F,G; Supplementary Figure 17D). In particular, the first bifurcation in the metagene hierarchy generates two metagenes corresponding to the two major branches in the developmental hierarchy^24^ (Figure 6A), one which includes Monocytes and Neutrophils, and another which includes Mast cells, Basophils, Megakaryocytes and Erythroid cells (Figure 6F,G). The second bifurcation splits the latter into two additional specific metagenes, one including Mast cells and Basophils, and another - Megakaryocytes and Erythroid cells (Figure 6F,G). The third bifurcation further splits the metagene corresponding to the Mast-Baso branch to two separate metagenes that are more specific to either Mast cells or Basophils. Similarly, the fourth bifurcation splits the metagene corresponding to the Monocyte-Neutrophil branch to two separate Monocyte and Neutrophil associated metagenes. Finally, the last bifurcations split the metagene corresponding to the Megakaryocyte-Erythroid branch to four metagenes distinguishing between Megakaryocytes and Erythroid cells.

In conclusion, bioIB metagenes characterize distinct biological processes linked to the underlying cellular populations, while the metagene hierarchy unveils the biological relationships interconnecting these populations.

## Discussion

We introduced bioIB, a scRNA-seq tailored framework for clustering genes with respect to a set of known cellular labels, based on the Information Bottleneck method. We have shown that bioIB metagenes, which are biologically interpretable, provide a meaningful compressed representation which exposes signal-specific molecular pathways underlying the biological variance between the cellular populations of interest. BioIB simultaneously extracts pathways associated with a specific label and exposes signal-associated gene programs, such as intermediate states, as shown in the context of AD neurons, and transition signatures, as demonstrated in the context of the EMT. Given single-cell data from human differentiating macrophages, with overlapping signals of organ-of-origin and developmental time, bioIB successfully extracted two distinct compressed data representations, each depicting the respective biological processes. bioIB also identified a subpopulation of disease-associated astrocytes in single-nucleus data from an AD mouse model, providing the genotype as the signal of interest. At last, we have shown that the metagene hierarchical structure, produced by the iterative application of the IB algorithm, exposes interconnections between metagenes and their respective cell types. We showcased this in the context of differentiating hematopoietic cells, where the bioIB hierarchical structure matched the expected developmental hierarchy of hematopoietic cell types.

BioIB stands out among available methods for supervised gene program discovery due to its ability to generate multiple informative gene signatures, associated with the specific cellular division of interest. This feature is particularly valuable for uncovering pathways linked to signal characteristics, such as intermediate states, transition signatures, and subpopulations with distinct phenotypes. Furthermore, as opposed to existing methods, bioIB can provide a hierarchical structure of the produced gene signatures, revealing the interconnections between the underlying cellular populations.

We conducted a comprehensive analysis of the framework’s robustness and stability, showing that bioIB metagenes remain highly consistent across random initializations, hyperparameter tuning, and under data perturbations, such as cell subsampling. Since bioIB is based on mutual information, its output is sensitive to the representation of each cell cluster in the data, both in terms of the cluster size and the number of enriched genes in it. That being said, we demonstrated that given a strong transcriptional signature differentiating the underrepresented cluster, bioIB remains robust to its signal, extracting the relevant gene programs despite class imbalance. Furthermore, while by design bioIB relies on input cellular labels, which might be a limitation when annotations are ambiguous, we show that when supervised with a small proportion of incorrect labels bioIB does not overfit and its output remains aligned with the true transcriptional signal.

As with a majority of computational methods, the bioIB output depends on a hyperparameter, β, controlling the level of compression. This is analogous to setting the number of clusters in a clustering algorithm, making this value data-specific. Here, the interpretability of the obtained metagenes allows the user to tune β to obtain the desired number of informative metagenes. We showed that the choice of β does not affect the structure of the compressed representation, such that the gene-to-metagene mapping at the corresponding compression levels remains highly stable. The current hierarchical bioIB formulation is limited in its scalability to data size, as it relies on the exact solution to the IB problem. This can be overcome, as we have done in this study, by focusing the analysis on highly informative genes. A natural extension to bioIB to overcome this limitation more generally in future work is using an existing variational IB solver which relies on neural approximation^55–57^.

In future work bioIB can be extended to extract multiple related data representations with respect to several variables of interest, based on the multivariate information bottleneck framework^58^. This paradigm might be particularly useful in analyzing gene expression data, allowing to simultaneously extract multiple encoded signals and analyze the corresponding biological processes. Furthermore, bioIB could be extended to produce signal-specific cell clusters, or metacells, retaining maximal possible information about a target gene subset, such as disease biomarkers.

Here we demonstrated that bioIB can provide efficient characterization of signals of interest encoded in single-cell data, such as cell type, disease state or organ-of-origin. BioIB can be generalized beyond single-cell gene expression data to additional types of biological data, such as bulk RNA-seq and proteomics data, to expose signal-specific optimally compressed representations. In summary, bioIB is expected to enrich biological data analysis by revealing the hierarchical, signal-specific structure encoded in complex datasets.

## Materials and methods

### The bioIB algorithm

The bioIB algorithm provides a compressed representation of scRNA-seq data with respect to a signal of interest. To do so it takes as input a cell (*N*) by gene (*G*) scRNA-seq measurements matrix, *D* ∈ *R* ^*N*×*G*^; following standard practice we suggest providing log-normalized counts as input. Additional input to bioIB is a vector of cell labels related to the signal of interest *S* ∈ *R*^*N*×1^, labeling every cell with one of *K* possible cell states of interest defined using *Y* = [1, …, *K*], such that *Y* = {*S*}. Given this input, the bioIB pipeline is composed of two main steps: (1) obtaining a probabilistic representation of the count matrix, and (2) using this representation as input for the Information Bottleneck (IB) algorithm.

#### 1. Obtaining a probabilistic data representation

We use the input count matrix *D* and signal of interest *S* to obtain the relevant probability distributions required for the IB algorithm; the conditional probability matrix of cell states given the genes *p*(*y*|*x*) and the gene probability vector *p*(*x*). To convert to probability space, we define the random variables of *C* ~*Cat*({*c*_1_, …, *c*_*N*_}), *X* ~*Cat*({*x*_1_, …, *x*_*G*_}), *Y* ~*Cat*({*y*_1_, …, *y*_*K*_}), respectively representing the *N* cells, *G* genes and *K* cell states of interest. The empirical distributions of these are then constructed using the input data (see Equations 1-3).

#### 2. The IB algorithm

The obtained probabilistic representations, Information Bottleneck (IB) algorithm. *p*(*y*|*x*) ∈ *R*^*K*×*G*^ and *p*(*x*) ∈ *R*^*G*^ are the input for the IB^3^ is a dimensionality reduction method, designed to extract the information from data *X* that is relevant for the prediction of another related variable *Y*, such that the choice of *Y* determines the relevant components of the signal encoded in *X*. Mutual information (MI) is used to evaluate both the extent of compression, 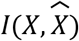, and the level of relevant information preserved in the compressed data, through 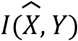. A trade-off parameter β is introduced to control the amount of compression (distortion) allowed. Formally, the IB objective is given by,

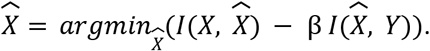

Notably, when β = 0, all genes are merged into one cluster (full compression), and when β = ∞, the compressed data is identical to the original full data, so every cluster is associated with one particular gene, 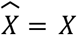. For every value of β, the algorithm yields the conditional probability matrix of *M* gene clusters, which we term metagenes, 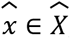, given the genes, 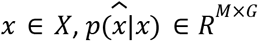, representing the optimal mapping of genes to metagenes, and the conditional probability matrix of cell states given the metagenes 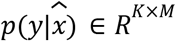. For the full mathematical description and the associated proofs for the information bottleneck algorithm, see refs^3,4^.

There are many ways to solve the IB objective (including neural approximators introduced recently^55–57^). Here we will focus on the Blahut-Arimoto algorithm^59^, described below. IB can provide either a series of solutions at different compression levels, using a reverse-annealing process, or a single solution with a flat division of the data points to a predefined number of clusters.

##### a. Blahut arimoto

*While True:*

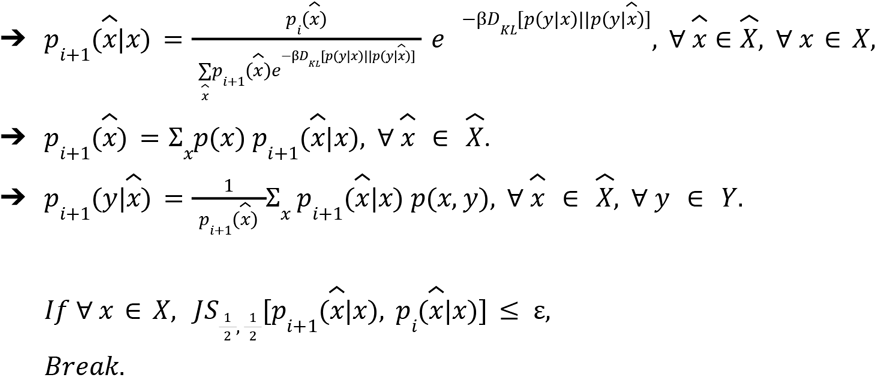

Here, ε is a threshold parameter used to define convergence based on the difference between previous and current iterations. For a given β, the algorithm converges into a stable solution, providing two output probability matrices that define 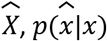 and 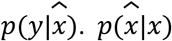 determines the mapping between the original data points *x* ∈ *X* to data clusters 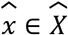, whereas 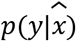 defines the association between the data clusters, 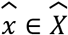, and the groupings of the signal of interest, *y* ∈ *Y*.

##### b. Flat Clustering

To achieve the division of the data points *x* ∈ *X* to a defined number of clusters *M*, we follow previous work^4^ and initialize the IB algorithm with a random mapping of *X* to *M* clusters, generating a binary conditional probability matrix 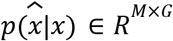. The corresponding 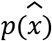 and 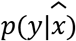 are obtained, using basic probability rules and Bayes Theorem. Since this process introduces a dependence of the output on the initialization, we randomly initialize the algorithm *n* = 100 times and select the mapping that minimizes the objective function (Eq.4).

##### c. Reverse-annealing

For the hierarchical mode of bioIB, in the process of reverse-annealing the IB algorithm is initialized with a compressed representation 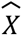 that is identical to the original data *X* and with a large value of β:

- 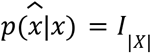
- 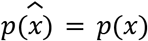
- 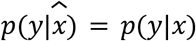
- β_*max*_ → ∞

Next, we run the algorithm iteratively, while reducing β. Upon convergence, we initialize the next iteration with the final 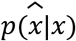 mapping achieved in the previous step, and with β − Δ, for a small step size Δ. Following this procedure we achieve a series of solutions for every value of β: ∀β ∈ {β_*min*_, β_*min*_ + Δ, …, β_*max*_}. At the end of this process β_*min*_ → 0, corresponding to maximal compression, where 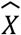 consists of a single point, uniting all the original data points in *X*. Reverse-annealing ultimately yields a hierarchical structure that mirrors several important aspects of the identified clusters, such as their informativity for discrimination between the labels of interest *Y*, as well as the interconnections among them. It is important to note that β controls the maximal number of metagenes, namely the number of end-nodes in the hierarchy, and modifying it does not affect the hierarchical structure itself, with consistent metagene-to-gene mapping at the corresponding hierarchy resolutions (Supplementary Figure 14). Furthermore, as opposed to flat bioIB clustering, hierarchical bioIB does not include a random initialization, and therefore its input is identical when consequently applied to the same data. While hierarchical bioIB is more computationally demanding than flat clustering, bioIB supports GPU acceleration for increased efficiency (Supplementary Figure 15, Supplementary Table 12).

### Downstream analyses

1. **Identifying representative genes**: The representative genes *x* ∈ *X* for a given metagene 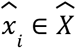 are identified as the ones that maximize 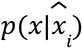. Specifically, for a given metagene, we first order the genes by their conditional probability 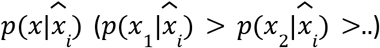. For a given *τ* ∈ [0, 1], the set of *j* representative genes {*x*_1_, *x*_2_, …, *x*_*j*_} is chosen as the minimal set such that:

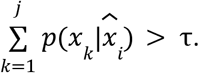
2. **Recovering single-cell metagene expression:** The bioIB output provides the mapping of the original count matrix *D* ∈ *R*^*N*×*G*^ to its compressed representation 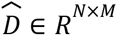. Namely, we obtain the weighted expression of genes, *x* ∈ *X*, using the mapping 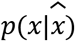., given by,

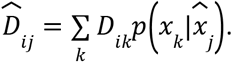 As a result, we obtain a cell (*N*) by metagene (*M*) compressed data matrix, 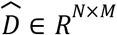, such that 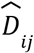 represents the expression level of metagene *j* in cell *i*.
3. **Clustering cells based on the relative metagene expression:** Based on single-cell metagene expression, each cell can be assigned to a metagene-associated cluster by identifying the metagene with the highest relative expression in that cell, given by:

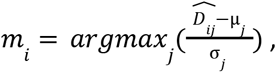 Where *m*_*i*_ is the metagene-associated cluster label of cell *i*, μ_*j*_ is the average expression of metagene *j* expression over all cells, and σ_*j*_ is the standard deviation of metagene *j* expression over all cells.
4. **Extracting the metagene hierarchy:** The bioIB reverse-annealing output provides a series of conditional probability matrices: 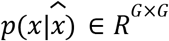 and 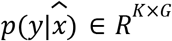 for each β. Since we initialize the reverse-annealing process with 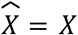, these matrices include *N* metagenes, but only *M* of them are unique. We first identify the most representative gene *x* of each metagene 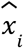, using 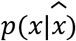:

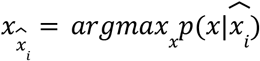 Next, we extract the metagene hierarchy by identifying the merging points of the most representative genes for each metagene across decreasing β. For example, metagenes 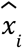 and 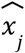 are considered merged at β_*merge*_ if 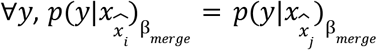. The identified merging points are recorded using a format of the scipy.cluster.hierarchy.linkage() output linkage matrix and plotted using scipy.cluster.hierarchy.dendrogram(). The code and the documentation for the relevant bioIB functions are provided in the bioIB package at https://github.com/nitzanlab/bioIB.
5. **Linking metagenes to cell types:** Metagenes, 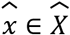, are linked to cell types, *y* ∈ *Y*, using 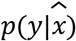 mapping, given by,

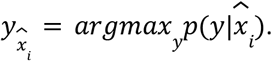 Metagene 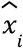 is identified as an intermediate metagene if its maximal probability is close to the uniform distribution, namely if 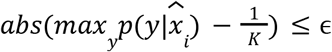, where ϵ stands for a similarity threshold (by default, ϵ = 0. 15).

### Datasets

#### NFT-free and NFT-bearing neurons from Alzheimer’s Disease (AD) human brains

##### Data preprocessing

We obtained the dataset of single-cell RNA-seq of NFT-free and NFT-bearing AD neurons from ref.^20^, available at https://cellxgene.cziscience.com/collections/b953c942-f5d8-434f-9da7-e726ba7c1481. We downloaded the dataset of excitatory cells and further filtered it to include only cells of Ex2 subtype, out of considerations of total cell number, similar frequencies of NFT-free and NFT-bearing neurons, and total number of differentially expressed genes^20^ resulting in 10,955 cells. Following basic preprocessing using scanpy’s sc.pp.normalize_per_cell() and sc.pp.log1p(), the data was further reduced to 3000 highly variable genes using scanpy’s sc.pp. highly_variable_genes() with the default parameters.

#### Method application

We applied bioIB to generate *m* metagenes using bioib.flat_clustering(*m*) with default parameters, for *m* = [2, 3], as for higher *m* additional metagenes showed no statistically significant enrichment in the gene set enrichment analysis. The gene set enrichment analysis was performed using gseapy’s enrichr() with GO_Biological_Process_2025 and KEGG_2021_Human gene sets.

#### Benchmarking

The ground-truth genes and pathways were obtained from the Supplementary Table 4 of ref.^20^ Out of all the identified pathways, we selected 150 pathways, enriched in the analyzed cluster Ex2, including 94 unique genes, appearing in the analyzed 3,000 top highly variable genes (the filtered genes and pathways are provided in Figure 2C and in Supplementary Table 4). The Proportion of recovered ground-truth genes was calculated as the proportion of top representative inferred genes that appear in the list of 94 genes representing the ground-truth pathways (Figure 2F).

We additionally benchmarked the correspondence between the gene signatures generated by different methods, and the ground-truth pathways. For this analysis, we used a deterministic version of gene signatures, where they were composed for each method by the top 500 inferred representative genes per signature, resulting in deterministic gene clusters of equal size. Pathway correspondence score *c*_*i*_ between pathway *i, P*_*i*_, and a set of gene signatures *M*_*j*_ ∈ *M*, was calculated as follows:

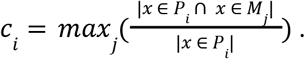

Thus, this score reflects the maximal proportion of genes in pathway *P*_*i*_ that represent the same gene signature *M*. For example, if the whole pathway is represented by the same signature, this pathway receives a maximal score of 1. We adjust for the underrepresented pathways, as follows: if |*x* ∈ *P*_*i*_ | ≤ 1, *c*_*i*_ = 0. We further aimed to correct the scores for the baseline correspondence, which depends on the total number of signatures and the total number of ground-truth genes, identified by different methods within the top 500 representative genes. For each pathway, we normalized each method’s performance by the mean correspondence score of the identified genes randomly assigned to the same number of signatures over 10 iterations (Figure 2G, Supplementary Figure 6).

We compared bioIB performance to scGeneFit, scANVI, NMF and LDVAE. We applied scGeneFit with default parameters (method=‘centers’, redundancy=0.25). We identified 600 total markers and then divided them into signatures based on the fold change between NFT-free and NFT-bearing neurons. For each signature, we inferred top representative genes based on expression fold change between different cellular groups clustered by input labels. scANVI was trained with default parameters, generating 10 latent representations in a fully supervised mode. The group-associated gene signatures were obtained with integrated gradients, as explained here - https://docs.scvi-tools.org/en/stable/tutorials/notebooks/use_cases/interpretability.html. ScanVI gene signatures were defined as top marker genes with the highest attribution means per cell group. NMF was applied with the default parameters using ‘sklearn.decomposition.NMF()’. The NMF gene signatures were defined as genes with top coefficients per factor, stored in nmf.components_. LDVAE was trained with the default parameters, generating *m* latent representations with *m* = 2, 3, 5, similarly to bioIB. The LDVAE gene signatures were obtained as genes with top loading scores per factor, assessed via model.get_loadings(), as explained here-https://docs.scvi-tools.org/en/stable/tutorials/notebooks/scrna/linear_decoder.html.

### EMT dataset from human mammary epithelial cells

#### Data preprocessing

We obtained the dataset of the human mammary epithelial cells across the epithelial-to-mesenchymal transition from ref.^21^, available at https://www.ncbi.nlm.nih.gov/geo/query/acc.cgi?acc=GSE114687. We further used the same preprocessing and annotation pipeline, as in ref^60^. The final analyzed dataset included 17,632 cells of HuMEC cell line and 5,000 top highly variable genes.

#### Method application

We applied bioIB to the obtained dataset to generate 3 metagenes using bioib.flat_clustering(3) with default parameters. The gene set enrichment analysis was performed using gseapy’s enrichr() with GO_Biological_Process_2025 and MSigDB_Hallmark_2020 gene sets, in alignment with the original study^21^.

### Multi-organ atlas of human differentiating macrophages

#### Data preprocessing

We obtained the dataset of the multi-organ atlas of human differentiating macrophages from ref.^22^, available at https://developmental.cellatlas.io/fetal-immune. We downloaded the dataset of myeloid cells and further filtered the data to include only cells of macrophage cell types. Using the gestational week label we assigned the cells into two groups, “Early” and “Late” (Early: < 14 weeks; Late: >= 14 weeks). In order to avoid bias towards less represented organ groups, we filtered out cells which originated from organs with less than 2800 total cells, resulting in cells originating from five organs: kidney (KI), liver (LI), skin (SK), spleen (SP) and yolk sac (YS). Following basic preprocessing for low-quality cells using scanpy’s^61^ ‘sc.pp.filter_cells(min_genes=200)’, the data used for bioIB analysis included 108,197 cells. We further reduced the data to 500 highly variable genes using scanpy’s^61^ ‘sc.pp.highly_variable_genes()’ with the default parameters.

#### Method application

We applied bioIB to the obtained dataset twice, (1) setting *Y* as the development stage (*Y* = [*Early, Late*]), and (2) setting *Y* as the organ-of-origin (*Y* = [*Kidney, Liver, Skin, Spleen, Yolk Sac*]). In both analyses we initialized the reverse-annealing process with β_*max*_ = 30. With *Y* set to be the development stage, bioIB initially produced five metagenes, with metagene 4 mostly represented by the marker genes of T-cells and B-cells and being enriched in only 229 out of 108,197 cells in the dataset (Supplementary Figure 13). Furthermore, we found that cells maximizing metagene 4 feature significantly higher doublet scores than cells maximizing the other four bioIB metagenes (Supplementary Figure 13). Therefore, we concluded that metagene 4 has identified a doublet subpopulation and excluded it from the downstream analysis.

### Astrocytes from a murine model of Alzheimer’s Disease (AD)

#### Data preprocessing

We obtained single-nucleus RNA-seq measurements from astrocytes from AD mouse model and wild-type (WT) mice from ref.^23^, available at https://www.ncbi.nlm.nih.gov/geo/query/acc.cgi?acc=GSE143758. Following normalization and log-transformation, we performed leiden clustering using Scanpy’s ‘sc.tl.leiden()’ function with default parameters. Following this, we retained the cell clusters with enriched expression of the astrocytic markers *Gfap* and *Slc1a3*, resulting in n=7036 cells. As a last step we extracted highly informative genes with respect to the signal of interest, or disease state, encoded by the provided genotype annotation *Y* = [*AD, WT*]. This was done by retaining the 1000 genes with the highest information gain (IG) values, where the IG is defined using the mutual information between the gene expression probability *p*(*x*) and the genotype probability *p*(*y*),

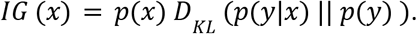

#### Method application

We applied bioIB with *Y* set as the mouse genotype: *Y* = [*AD, WT*]. The reverse-annealing process was initialized with β_*max*_ = 150.

#### Constructing a list of AD-related genes

AD-associated genes were defined as differentially expressed genes in at least 7 of the 15 AD-APP mouse model studies as part of the AD meta-analysis resource, which has summarized and compared the differential expression results from a wide range of AD transcriptomic studies^48^.

### Hematopoiesis dataset

#### Data preprocessing

We obtained the dataset of the differentiating hematopoietic cell types collected by ref.^24^ and processed by ref.^49^. Data was downloaded using the Cospar package (https://cospar.readthedocs.io/en/latest/index.html) using the function ‘cs.datasets.hematopoiesis()’). We filtered out the undifferentiated cells, as well as the differentiated cell types with less than 300 total cells, resulting in a data subset of 27387 cells. We further reduced the data to the highly variable genes using scanpy’s^61^ ‘sc.pp.highly_variable_genes()’ with the default parameters, resulting in 1803 genes.

#### Method application

We calculated the IG values (as above) for the highly variable genes and used as input for bioIB the 300 genes with the highest IG values.

### Simulated Data

For bioIB evaluation and benchmarking, we applied it to simulated datasets, generated using Splatter^25^. The cellular division to groups of interest was done using method=‘groups’. The simulation parameters are provided in figure captions of the corresponding Supplementary Figures (1-4).

## Data availability

The datasets analyzed in the current study are available at:

- Immune macrophage atlas: https://developmental.cellatlas.io/fetal-immune
- Alzheimer’s Disease astrocytes: https://www.ncbi.nlm.nih.gov/geo/query/acc.cgi?acc=GSE143758
- Hematopoiesis: https://cospar.readthedocs.io/en/latest/index.html

## Code availability

Software is available at https://github.com/nitzanlab/bioIB.

## Acknowledgements

We would like to thank the late Professor Naftali Tishby for initiating this project and his guidance which made this work possible. We would also like to express our gratitude to Professor Eli Nelken, Hadar Levi Aharoni, and Shlomi Agmon for fruitful discussions. We acknowledge all members of the Nitzan lab for general feedback.

This work was supported by the Azrieli, Kaete-Klausner and TEVA PhD fellowships (S.D.), a scholarship for outstanding doctoral students in data-science by the Israeli Council for Higher Education and the Clore Scholarship for PhD students (Z.P.), an Alon Fellowship, the Israel Science Foundation (Grant no. 1079/21), and the European Union (ERC, DecodeSC, 101040660) (M.N.). Views and opinions expressed are however those of the author(s) only and do not necessarily reflect those of the European Union or the European Research Council.

