## Supplementary Figures for "Identifying maximally informative signal-aware representations of single-cell data using the Information Bottleneck"

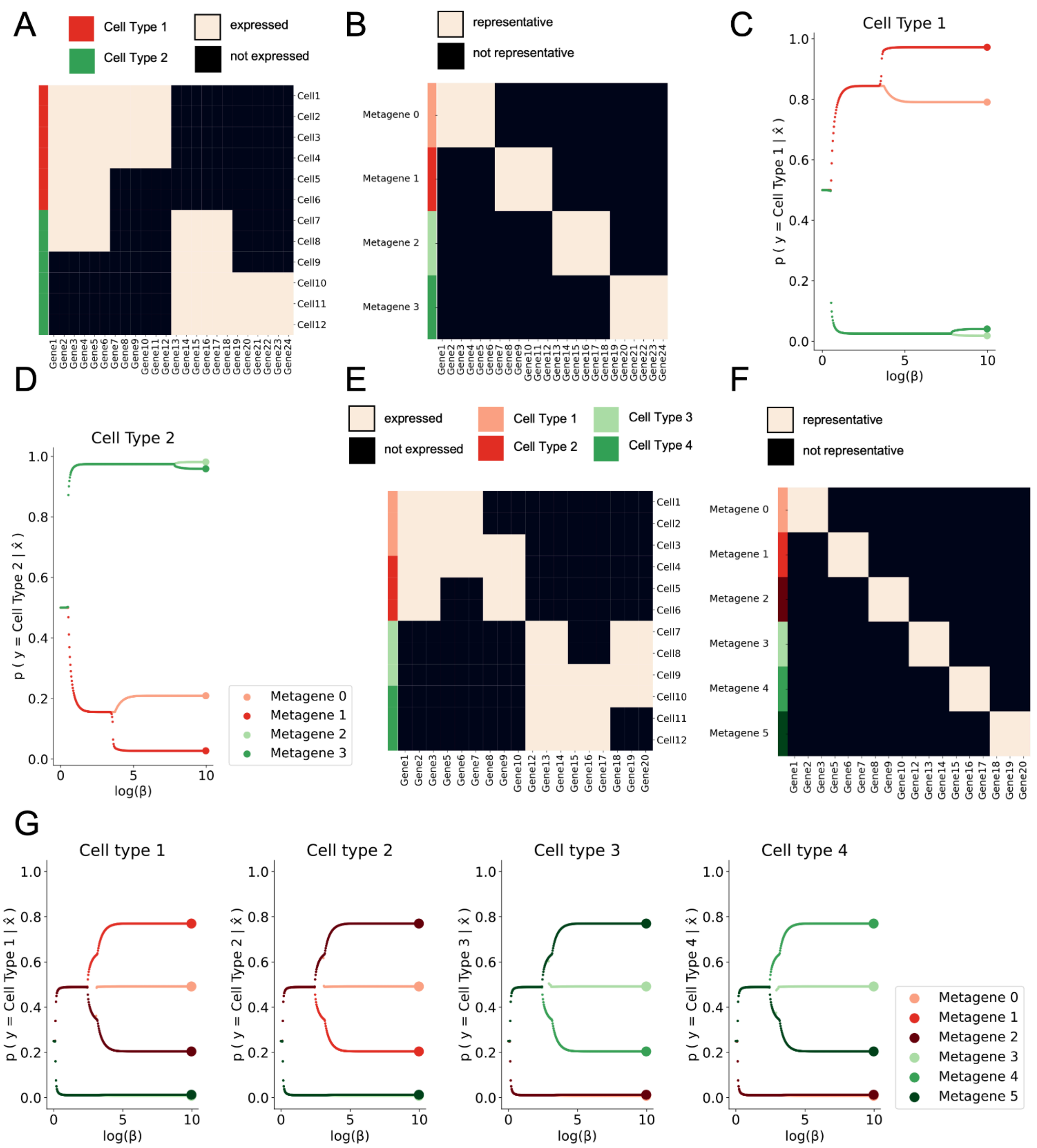


**Supplementary Figure 1. Hierarchical bioIB-enabled downstream analysis over the toy datasets**

A) Heatmap representing the binary toy dataset X (white - expression, black - no expression). A total of 12 cells are categorized to two cell types, six cells in each. B) Heatmap representing the $p(x|\hat{x})$ probability matrix, showing the genes associated with each metagene. For example, genes 1-6 belong to metagene 0, genes 7-12 belong to metagene 1, etc. C) bioIB reverse annealing output given the input data shown in A. The panel shows the metagenes with respect

to their prediction value for cell type 1, $p(y = Cell\,Type\ 1|\hat{x})$. The order of metagene bifurcations reflects their importance in differentiating between cell types. The metagenes are first divided into two branches, corresponding to their respective association with the two cell types: metagenes 0 and 1 are enriched in cell type 1, whereas metagenes 2 and 3 are enriched in cell type 2. The second bifurcation separates the branch linked to cell type 1 into a metagene that is uniquely expressed in cell type 1 and metagene 2 that also exhibits expression in cell type 2, whereas the last bifurcation distinguishes between two metagenes which are both uniquely expressed in cell type 2. D) Same as C with respect to cell type 2.
E-H) Analysis of a binary synthetic count matrix (white - expression, black - no expression). 12 cells are categorized to four linked cell types, cell types 1 and 2 that share a transcriptional program, and cell types 3 and 4, also sharing a program. E) Heatmap representing the dataset. F) Heatmap of the probability matrix, $p(x|\hat{x})$, showing the genes associated with each metagene. H) bioIB reverse annealing output of the toy dataset presented in E. Panels from left to right present the metagenes with respect to their prediction value for cell types 1-4.

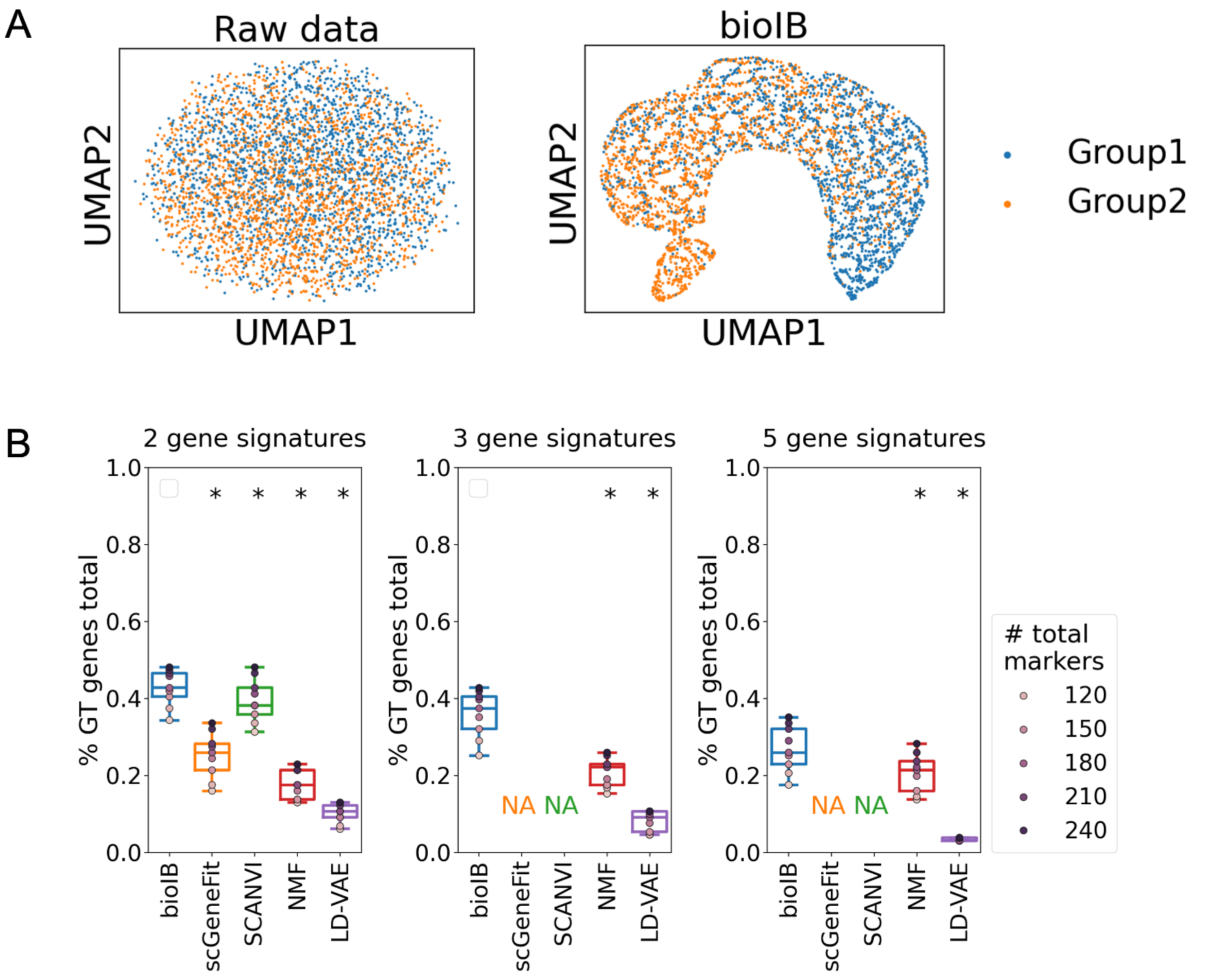


**Supplementary Figure 2. Benchmarking flat bioIB clustering on simulated data.**
A) UMAP representing the Splatter[1]-simulated dataset before (left) and after (right) bioIB processing. The simulation parameters were batchCells=4000, nGenes=4000, de.prob=0.03, de.facScale=0.03. The analysis was performed on 3000 top highly variable genes. The bioIB-compressed data consists of 4000 cells and two group-specific metagenes. B) Benchmarking of the proportion of ground-truth genes (Y axis) captured within 100-300 top inferred representative genes exposed by different methods using 2,3 and 5 gene signatures (Methods). Ground-truth genes were defined by the Splatter rowData output. Every gene with 'DEFac' parameter smaller or larger than 1 was considered a ground-truth relevant gene. In box plots middle line, median; box boundary, interquartile range (IQR); whiskers, 1.5*IQR; gray dots, points beyond the minimum or maximum whisker. Statistical significance was assessed using the WIlcoxon signed rank test (nonparametric), with * indicating $p < 0.01$, in comparison to the bioIB scores.

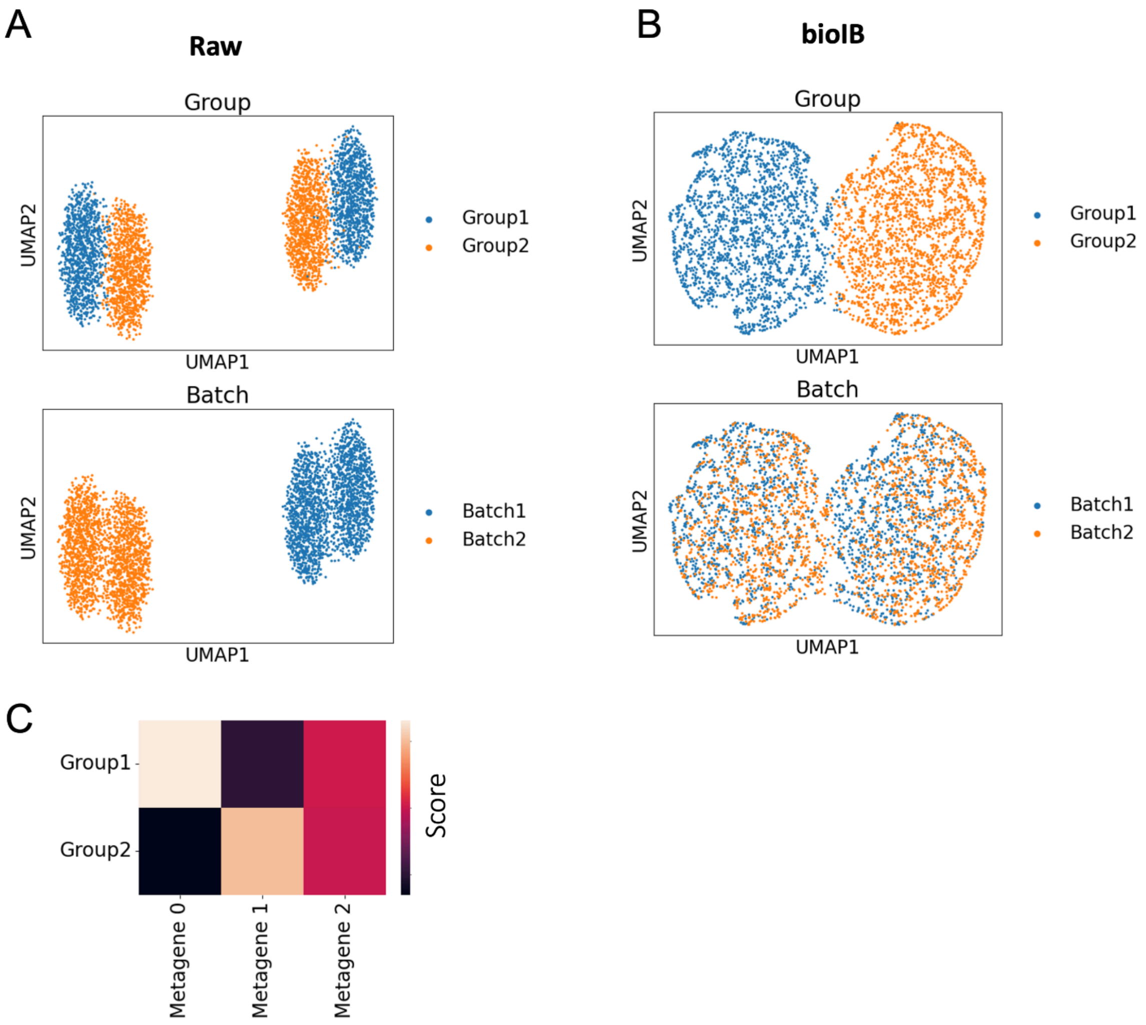


**Supplementary Figure 3. bioIB performs well despite batch effects.**
A) UMAP representing the Splatter[1]-simulated dataset, colored by group, representing the signal of interest (top) and batch (bottom). The simulation parameters were batchCells=(2000,2000), ngenes=3000, de.prob=0.01, de.facLoc=0.05. The analysis was performed on 3000 top highly variable genes. B) UMAP representing the bioIB-compressed data of 4000 cells by three metagenes, generated with the signal of interest set to Group, colored by group, representing the signal of interest (top), and batch (bottom). C) Heatmap representing the three bioIB metagenes and the corresponding metagene-to-cell state probabilistic mapping score ($p(y|\hat{x})$).

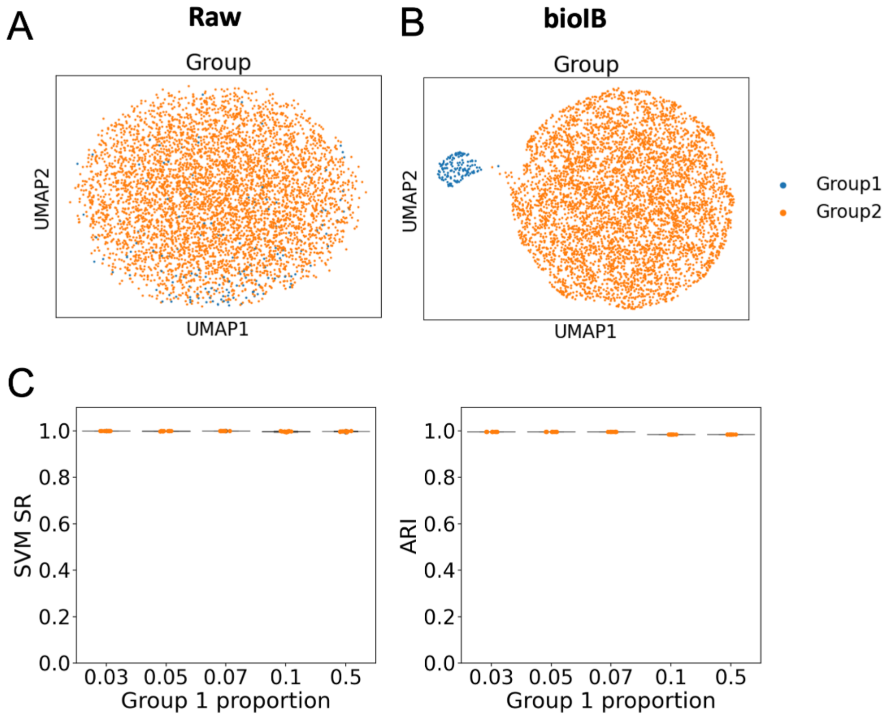


**Supplementary Figure 4. bioIB performs well in the context of class imbalance.**
A) UMAP representing the Splatter[1]-simulated dataset, mimicking a context of class imbalance, with only 3% of cells labeled as Group 1, and 97% of cells as Group 2. The simulation parameters were batchCells=5000, nGenes=6000, de.prob=0.1, de.facScale=0.1. The analysis was performed on 3000 top highly variable genes. B) UMAP representing the bioIB-compressed data of 5000 cells by three metagenes, generated with the signal of interest set to Group. C) Benchmarking the accuracy of bioIB metagenes in predicting cellular labels (Group 1 vs Group 2), in the context of class imbalance of 3, 5, 7, 10, and 50%, representing a regular balanced dataset. Left: Success rate of support vector machine (SVM) classifier in predicting the cellular class, while trained on bioIB metagenes. The experiment was repeated n=10 times with random data divisions (test size=0.2). Right: Adjusted Rand Index between two cell clusters generated using agglomerative clustering based on bioIB metagenes, and the ground-truth division of cells to Group 1 and Group 2. In box plots middle line, median; box boundary, interquartile range (IQR); whiskers, 1.5*IQR; gray dots, points beyond the minimum or maximum whisker.

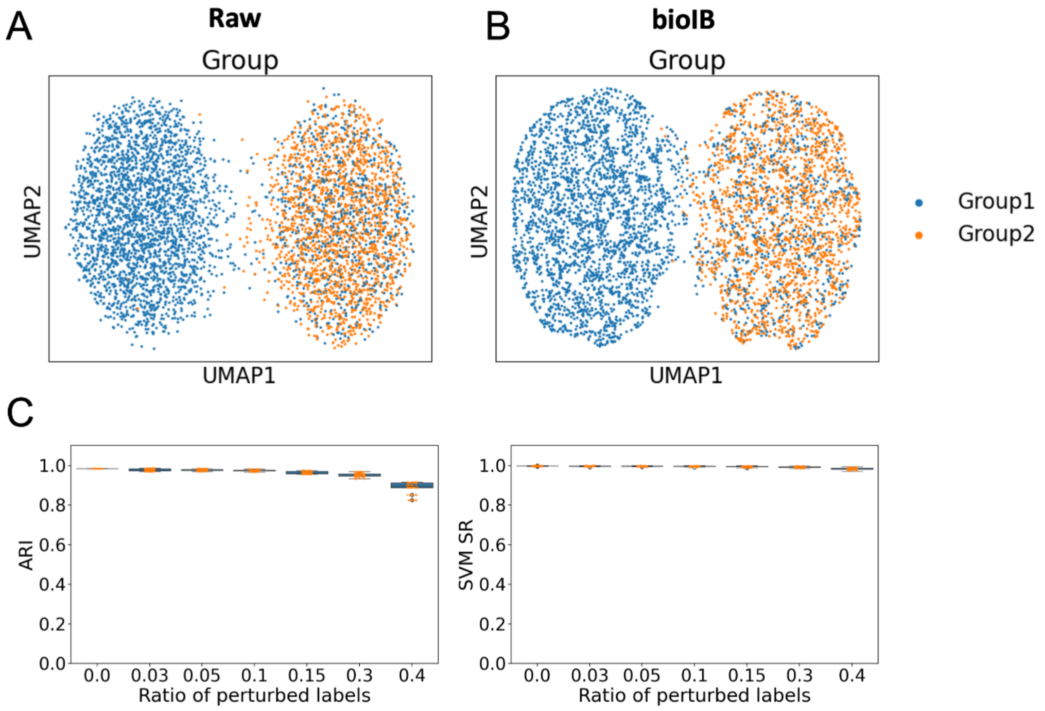


**Supplementary Figure 5. bioIB performs well in the context of erroneous cell annotation.** A) UMAP representing the Splatter[1]-simulated dataset, mimicking a context of erroneous cell annotation, with 15% of cells in Group 2 erroneously labeled as Group 1. The simulation parameters were batchCells=5000, nGenes=6000, de.prob=0.1, de.facScale=0.1. The analysis was performed on 5000 top highly variable genes. B) UMAP representing the bioIB-compressed data of 5000 cells by three metagenes, generated with the signal of interest set to Group. The erroneously labeled cells are clustered together with their true group (Group 2). C) Benchmarking the accuracy of bioIB metagenes in predicting the correct cellular labels (Group 1 vs Group 2), while trained on data with the erroneous annotation of 0, 3, 5, 10, 15, 30 and 40% of the cells. Left: Adjusted Rand Index between two cell clusters generated using agglomerative clustering based on bioIB metagenes, and the ground-truth division of cells to Group 1 and Group 2. Right: Success rate of support vector machine (SVM) classifier in predicting the cellular class, while trained on bioIB metagenes. The experiment was repeated n=10 times with random data divisions (test size=0.2). In box plots middle line, median; box boundary, interquartile range (IQR); whiskers, 1.5*IQR; gray dots, points beyond the minimum or maximum whisker.

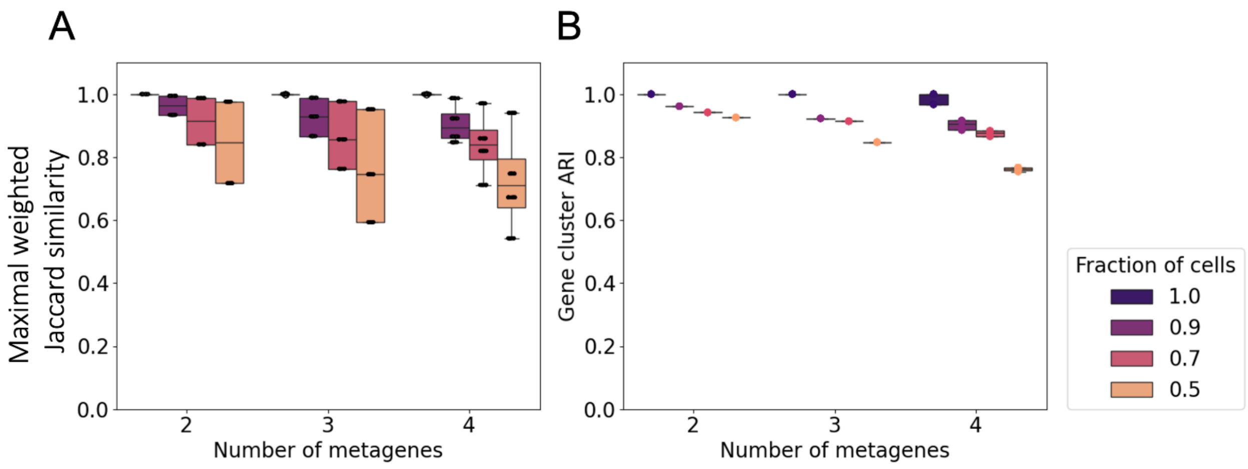


**Supplementary Figure 6. bioIB metagenes remain stable when the data is slightly perturbed.**
Comparison between the bioIB metagenes produced based on Splatter[1]-simulated data with two cell groups, provided as labels, in the context of cell subsampling. The simulation parameters were batchCells=5000, nGenes=6000, de.prob=0.1, de.facScale=0.1. The analysis was performed on 5000 top highly variable genes. The cells were subsampled, with keeping 1.0 (control), 0.9, 0.7 and 0.5 fractions of cells, stratified by the signal of interest ('Group' label). We compared both the probabilistic mappings between genes and metagenes, using the maximal weighted Jaccard similarity (A; Supplementary Methods) and binary assignment of genes to metagene-derived clusters using the Adjusted Rand Score (B; Supplementary Methods). The cluster is defined as top 50 representative genes per metagene. In box plots middle line, median; box boundary, interquartile range (IQR); whiskers, 1.5*IQR; gray dots, points beyond the minimum or maximum whisker.

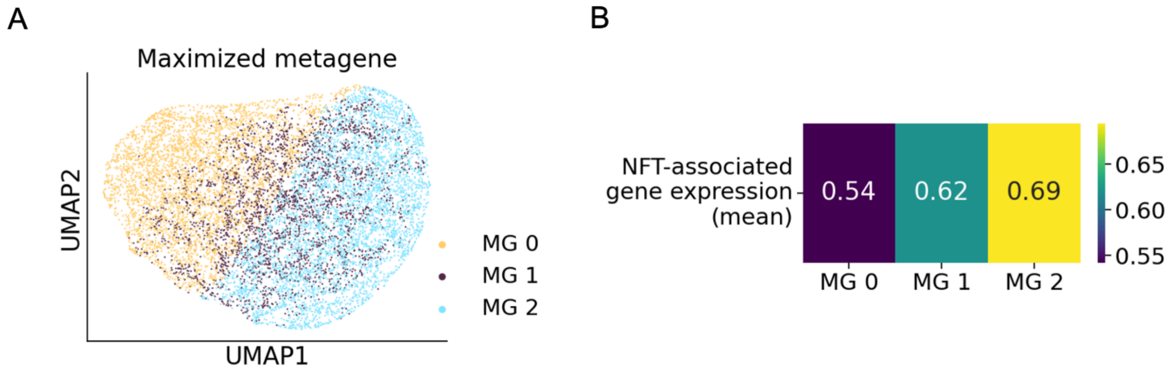


**Supplementary Figure 7. NFT-associated gene markers are expressed at an intermediate level in bioIB metagene 1.**

A)UMAP representation of the bioIB compressed data from ref.[2], colored by the relatively maximized metagene (Methods). B) Heatmap showing the mean log-normalized expression of 94 NFT-associated gene markers, as shown in Main Figure 2C, in cell groups divided by the maximized metagenes, as shown in A.

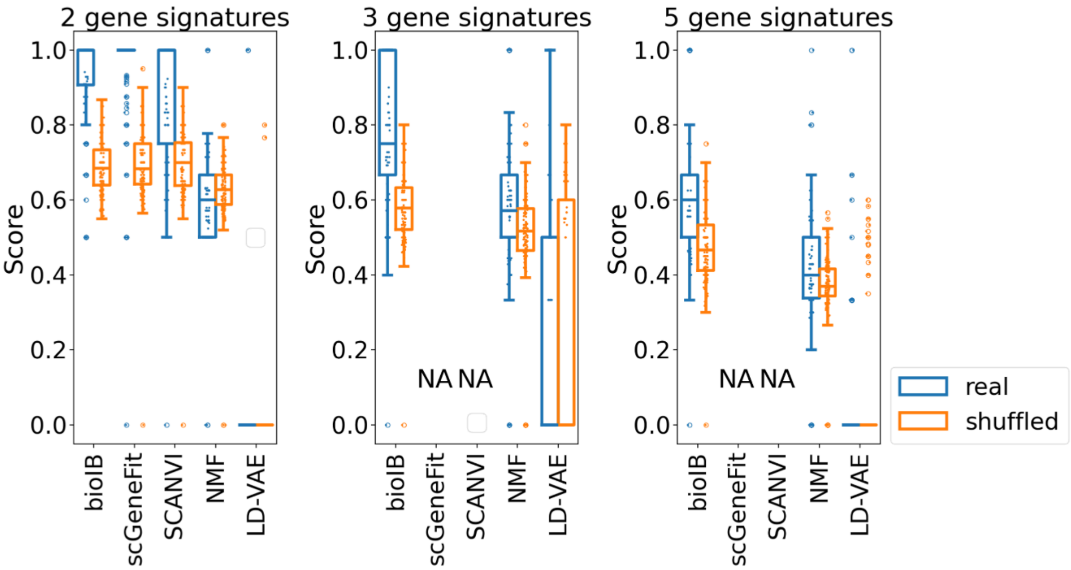


**Supplementary Figure 8. Benchmarking the correspondence between the gene signatures and the biological pathways in the data from NFT-free and NFT-bearing AD neurons.**

We evaluated the correspondence between the ground-truth biological pathways and gene signatures, produced by different methods, using the scRNA-seq dataset from NFT-free and NFT-bearing neurons[2]. For each pathway, we calculated the proportion of genes clustered together in the same signature by different methods, representing the biological coherence of the corresponding gene signatures (Methods). To account for differences in the number of ground-truth genes captured by each method, we calculated the same score for a shuffled control: genes captured by each method were randomly reassigned to gene signatures of the same sizes as the real ones. This shuffling was repeated $n = 10$ times. For each pathway, the real score is shown in blue, and the average score from the shuffled controls is shown in orange. In box plots middle line, median; box boundary, interquartile range (IQR); whiskers, 1.5*IQR; gray dots, points beyond the minimum or maximum whisker.

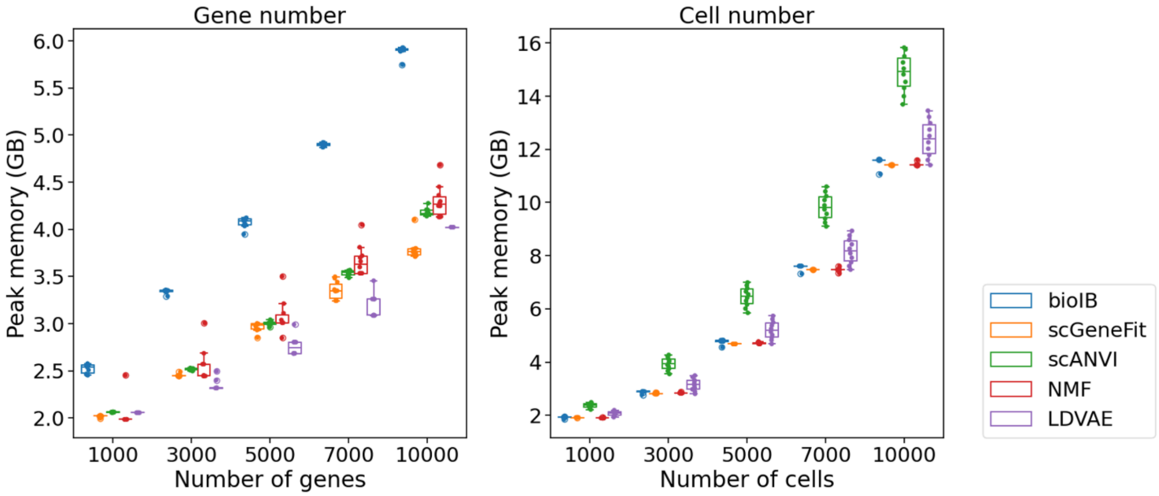


**Supplementary Figure 9. Comparing the CPU memory usage of bioIB to that of competing methods.**

(Left) CPU Runtime vs. number of genes given 10,955 cells. (Right) Runtime vs. number of cells given 3,000 highly variable genes for bioIB, scGeneFit, scANVI, NMF and LDVAE. The experiment was repeated $n = 10$ times.

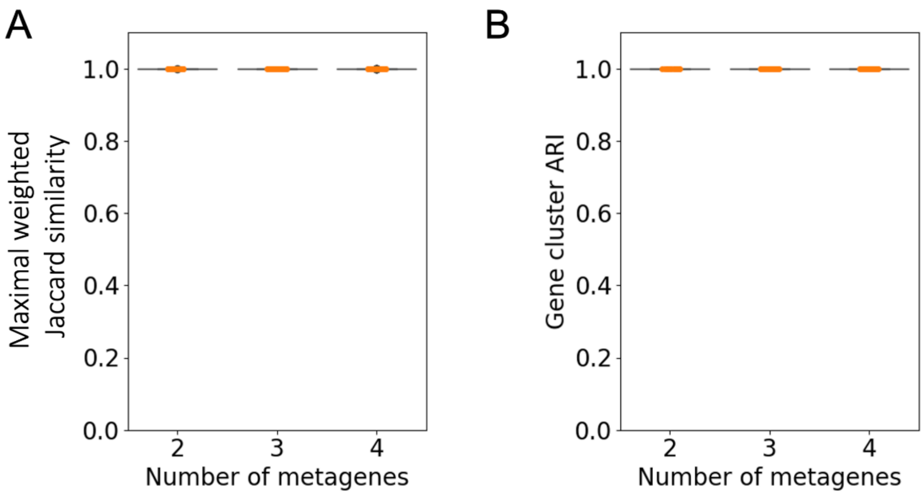


**Supplementary Figure 10. bioIB metagenes are stable across different method initializations.**
Comparison between the bioIB metagenes produced as a result of the consecutive application of flat bioIB clustering to the scRNA-seq dataset from NFT-free and NFT-bearing AD neurons[2]. The experiment was repeated $n = 10$ times. We compared both the probabilistic mappings between genes and metagenes, using the maximal weighted Jaccard similarity (A; Supplementary Methods) and the binary assignment of genes to metagene-derived clusters using the Adjusted Rand Score (B; Methods). The cluster is defined as top 50 representative genes per metagene. In box plots middle line, median; box boundary, interquartile range (IQR); whiskers, 1.5*IQR; gray dots, points beyond the minimum or maximum whisker.

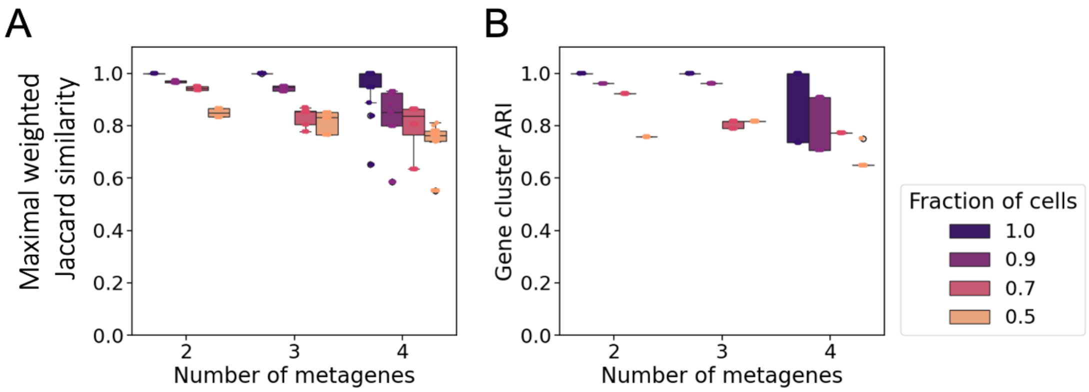


**Supplementary Figure 11. bioIB metagenes remain stable when the data is slightly perturbed.**

Comparison between the bioIB metagenes produced based on samples from the scRNA-seq dataset from NFT-free and NFT-bearing AD neurons[2]. The cells were subsampled, with keeping 1.0 (control), 0.9, 0.7 and 0.5 fractions of cells, stratified by the signal of interest (the presence of pathology). We compared both the probabilistic mappings between genes and metagenes, using the maximal weighted Jaccard similarity (A; Supplementary Methods) and binary assignment of genes to metagene-derived clusters using the Adjusted Rand Score (B; Supplementary Methods). The cluster is defined as top 50 representative genes per metagene. In box plots middle line, median; box boundary, interquartile range (IQR); whiskers, 1.5*IQR; gray dots, points beyond the minimum or maximum whisker.

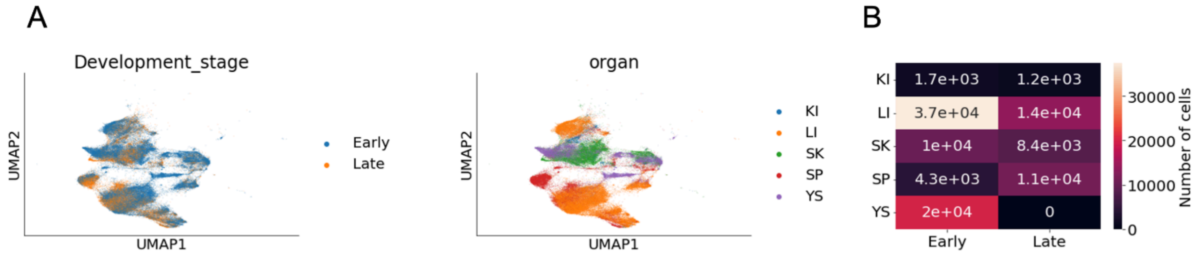


**Supplementary Figure 12. Visualizations and measures of developing macrophages.**
A) UMAP visualization of the analyzed scRNA-seq data[3] of developing macrophages, colored by developmental stage (left) and organ-of-origin (right). B) The number of cells in each combination of organ-of-origin (rows) and development stage (columns). Early: Early developmental stage (<14 weeks); Late: Late developmental stage (>=14 weeks); KI: kidney; LI: liver; SK: skin; SP: spleen; YS: yolk sac.

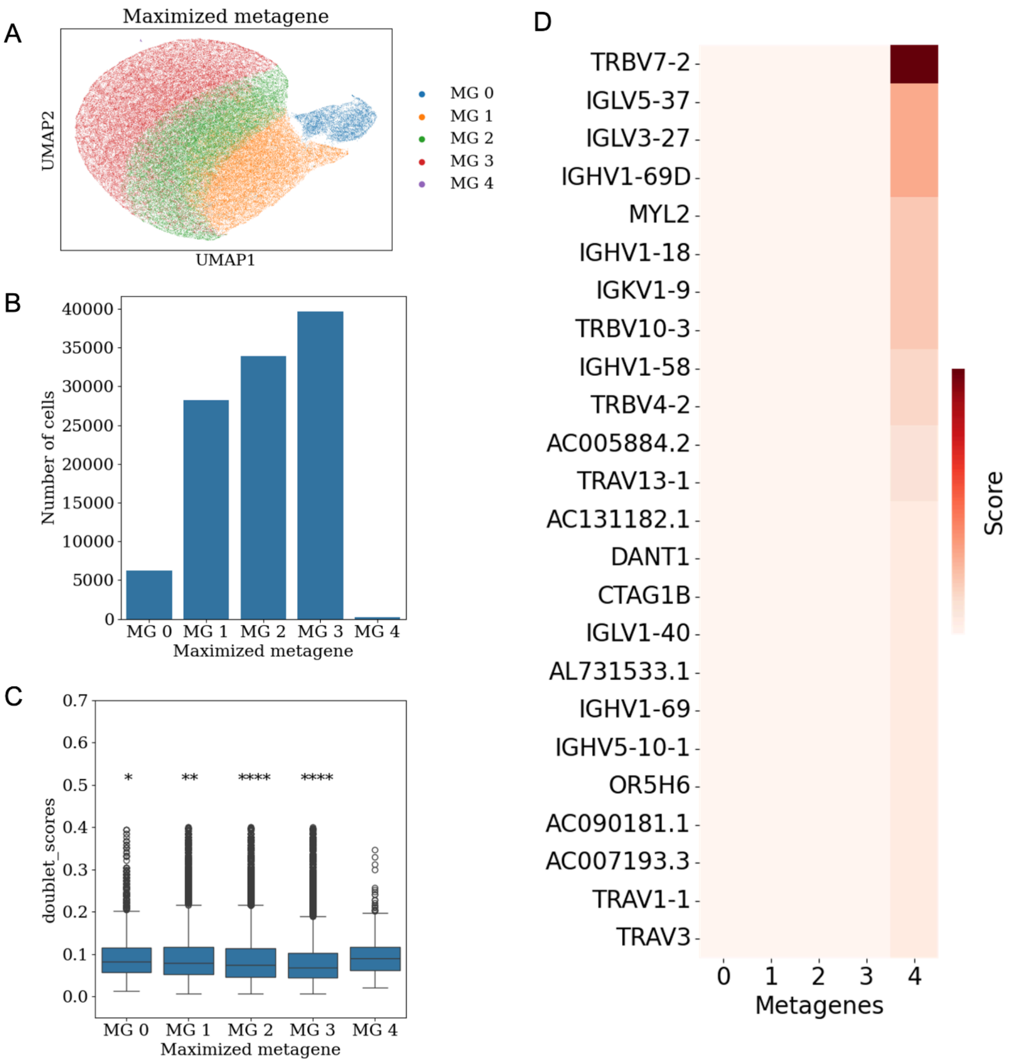


**Supplementary Figure 13. Characterizing bioIB metagenes, derived from the scRNA-seq data of developing macrophages, with $Y$ set to developmental stage.**

A)UMAP representation of single-cell RNA-seq data of developing macrophages[3], following bioIB compression with $Y$ set to developmental stage ('Early' or 'Late'). The colors represent metagene-associated cell clusters, with each cluster maximizing the corresponding metagene (Methods). B) Barplot showing the numbers of cells in each cluster shown in A. C) Boxplot showing the cellular doublet scores (provided by ref.[3]), divided by clusters shown in A. Statistical significance was assessed using the Mann–Whitney U-test (nonparametric), with **** standing for $p < 0.0001$, *** for $p < 0.001$, ** for $p < 0.01$, * for $p < 0.05$ and no symbol for non-significant results. D) Heatmap showing all the genes (with $p(x|\hat{x}) > 0$) representing metagene 4.

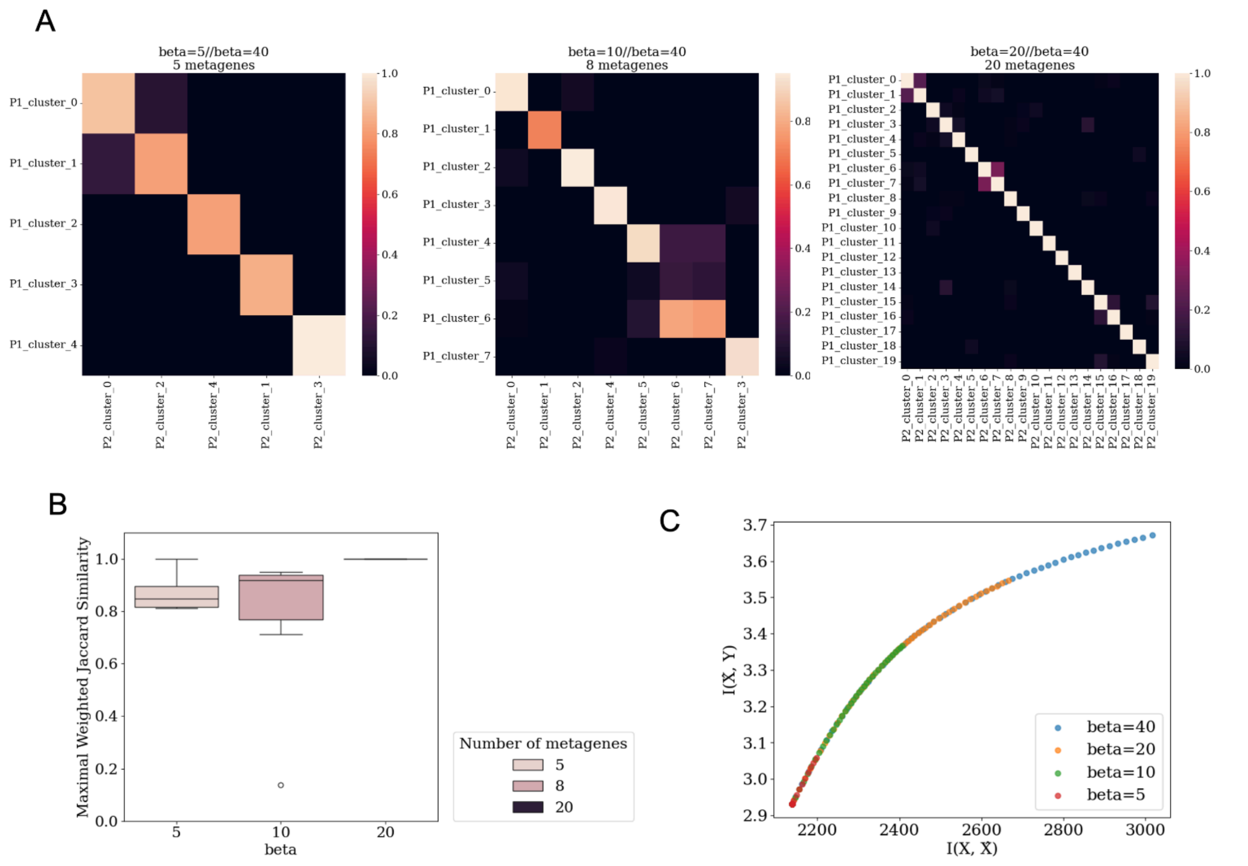


**Supplementary Figure 14. Stability and robustness of the hierarchical bioIB applied to the scRNA-seq dataset from developing macrophages[3].**

A)Heatmaps representing the weighted Jaccard similarity (Methods) between the bioIB metagenes generated given the initial $\beta = 40$ with the metagenes generated given the initial $\beta = 5$ (left); $\beta = 10$ (middle) and $\beta = 20$ (right), at the corresponding hierarchy resolutions (yielding equal metagene numbers). B) Maximal weighted Jaccard similarity per metagene in the bioIB compressed representation initiated with $\beta = 40$, compared with compressed data representations initiated with $\beta = 5,\ \beta = 10,\ \beta = 20$, at the corresponding hierarchy resolutions (5, 8 and 20 metagenes, respectively). C) The compression-information trade-off, showing the relevant information $I(\widehat{X}, Y)$ as a function of compression $I(X,\ \widehat{X})$ for hierarchical bioIB representations initialized by different $\beta$ values. In box plots middle line, median; box boundary, interquartile range (IQR); whiskers, 1.5*IQR; gray dots, points beyond the minimum or maximum whisker.

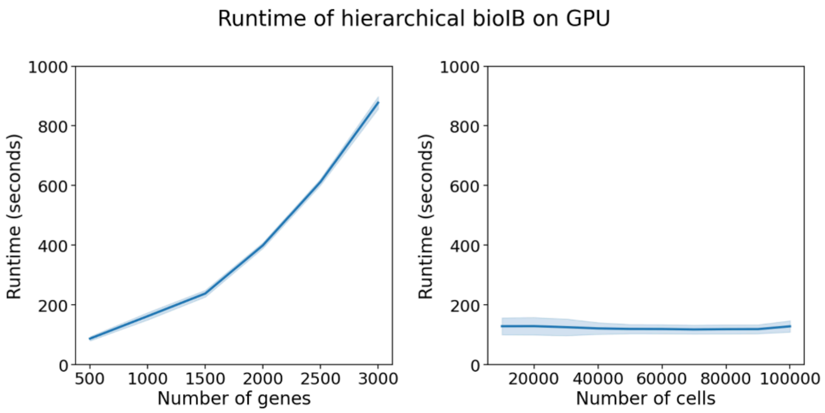


**Supplementary Figure 15. GPU-accelerated runtime of hierarchical bioIB, based on the scRNA-seq dataset from developing macrophages[3].**

Runtime (seconds) as a function of the number of highly variable genes (left) with 108,197 cells, or cells with 500 highly variable genes (right) in a GPU-accelerated analysis of hierarchical bioIB.

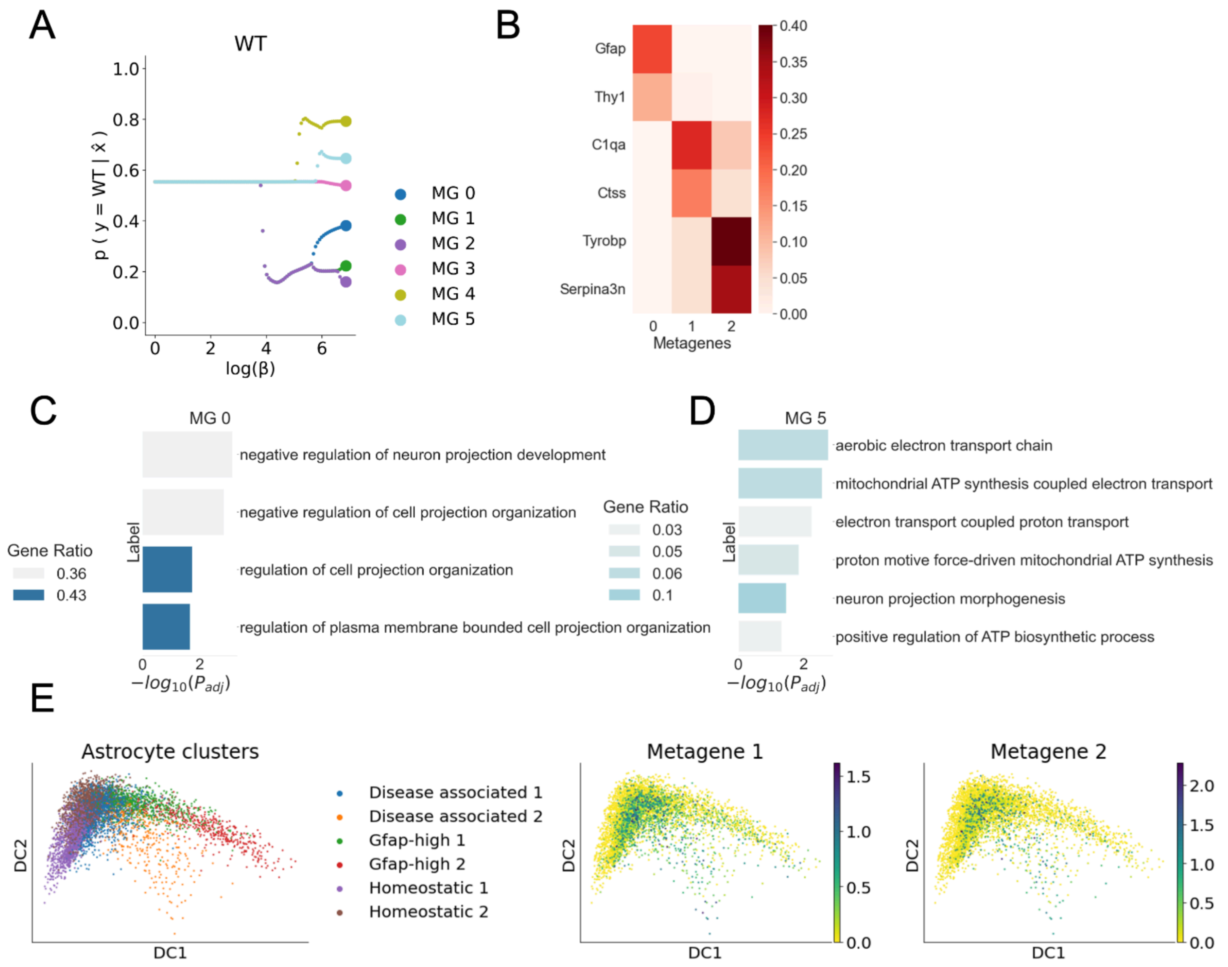


**Supplementary Figure 16. Studying Alzheimer disease-associated astrocytes using bioIB.** A) BioIB metagene hierarchy produced given the preprocessed snRNA-seq data[4], relating to the WT group. B) Heatmap showing p(x|x) values of the top representative genes for the metagenes 0,1,2. C, D) Gene Ontology biological processes significantly enriched among genes representative of metagene 0 (C) and 5 (D) in the bioIB hierarchy in (A) and in Figure 3A. Metagene 5, which is WT-related, is enriched with mitochondrial ATP synthesis, which was shown to be dysfunctional in AD[4]. E) Diffusion map embedding colored by the 6 clusters identified by ref.[4] (left), expression of bioIB metagene 1 (middle) and bioIB metagene 2 (right) over the representation on the left.

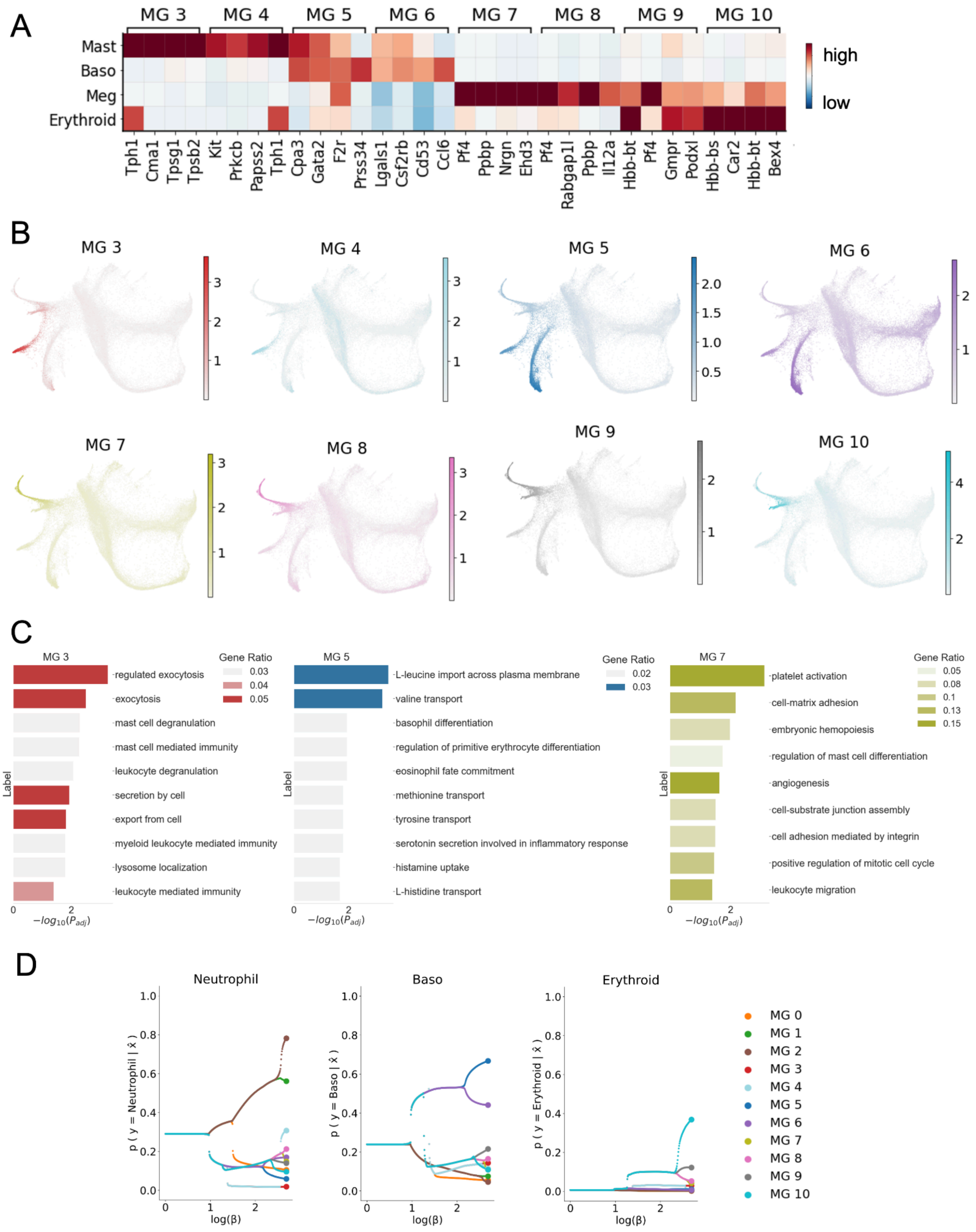
A
MG 3
MG 4
MG 5
MG 6
MG 7
MG 8
MG 9
MG 10
Mast
Baso
Meg
Erythroid
high
low
Tph1 Cma1 Tpsg1 Tpsb2 Kit Prkcb Papss2 Tph1 Cpa3 Gata2 F2r Prss34 Lgals1 Csf2rb Cd53 Ccl6 Pf4 Ppbp Nrgn Ehd3 Pf4 Rabgap1l Ppbp Il12a Hbb-bt Pf4 Gmpr Podxl Hbb-bs Car2 Hbb-bt Bex4
B
MG 3
MG 4
MG 5
MG 6
MG 7
MG 8
MG 9
MG 10
C
MG 3
Gene Ratio
0.03
0.04
0.05
regulated exocytosis
exocytosis
mast cell degranulation
mast cell mediated immunity
leukocyte degranulation
secretion by cell
export from cell
myeloid leukocyte mediated immunity
lysosome localization
leukocyte mediated immunity
Label
$-log_{10}(P_{adj})$
MG 5
Gene Ratio
0.02
0.03
L-leucine import across plasma membrane
valine transport
basophil differentiation
regulation of primitive erythrocyte differentiation
eosinophil fate commitment
methionine transport
tyrosine transport
serotonin secretion involved in inflammatory response
histamine uptake
L-histidine transport
MG 7
Gene Ratio
0.05
0.08
0.1
0.13
0.15
platelet activation
cell-matrix adhesion
embryonic hemopoiesis
regulation of mast cell differentiation
angiogenesis
cell-substrate junction assembly
cell adhesion mediated by integrin
positive regulation of mitotic cell cycle
leukocyte migration
D
Neutrophil
Baso
Erythroid
p ( y = Neutrophil | x̂ )
p ( y = Baso | x̂ )
p ( y = Erythroid | x̂ )
log(β)
MG 0
MG 1
MG 2
MG 3
MG 4
MG 5
MG 6
MG 7
MG 8
MG 9
MG 10

**Supplementary Figure 17. BiolB hierarchical representation reveals developmental connections between hematopoietic cell types.**
A)Heatmap showing scaled expression of the top representative genes of metagenes 3-10 (as in Figure 4D) across mast cells, basophils, megakaryocytes and erythroids. B) SPRING visualizations of the dataset[5], adapted from ref.[6] colored by the expression of metagenes 3-10. *MG – metagene. C) Gene Ontology enrichment results showing biological process categories significantly enriched in metagene 3 (left), 5 (middle) and 7 (right). D) Bifurcation plots of further compression of the 11 metagenes shown in Figure 4B relative to Neutrophils, Basophils and Erythroids.

**Supplementary Methods**

Metagene comparison metrics

1. Maximal weighted Jaccard similarity

To compare the metagene-to-gene probabilistic mappings $p(x|\hat{x})$ between different metagenes, we calculated the weighted Jaccard similarity, given by:

$$J(\hat{x}_i, \hat{x}_j) = \frac{\sum_x min(p(x|\hat{x}_i), p(x|\hat{x}_j))}{\sum_x max(p(x|\hat{x}_i), p(x|\hat{x}_j))}$$

Next, to identify the matched metagenes between different compressed representations, we identified the maximal weighted Jaccard similarity for every metagene, given by:

$$J_{max}(\hat{x}_i) = max_j J(\hat{x}_i, \hat{x}_j)$$

2. Adjusted Rand Index

To compare the bioIB compressed representations as systems of deterministic gene clusters, we first generated deterministic gene clusters by selecting $n$ top representative genes $x$ for each metagene $\hat{x}$, that maximize $p(x|\hat{x})$ (Main Methods). Next, we computed the Adjusted Rand Index[7] between the resulting deterministic gene clusterings, using the scikit-learn adjusted_rand_score(). Genes not assigned to any cluster in one of the clusterings were labeled with a placeholder and included in the comparison.
